# The deadly touch: protein denaturation at the water-air interface and how to prevent it

**DOI:** 10.1101/400432

**Authors:** Edoardo D’Imprima, Davide Floris, Mirko Joppe, Ricardo Sánchez, Martin Grininger, Werner Kühlbrandt

**Affiliations:** Max Planck Institute of Biophysics, Department of Structural Biology, Frankfurt am Main, Max von Laue Strasse 3, 60438, Germany; Buchmann Institute for Molecular Life Sciences, Goethe University Frankfurt, Institute of Organic Chemistry and Chemical Biology, Frankfurt am Main, Max von Laue Strasse 15, 60438, Germany; Sofja Kovalevskaja Group, Max Planck Institute of Biophysics, Frankfurt am Main, Max von Laue Strasse 3, 60438, Germany

## Abstract

Electron cryo-microscopy analyzes the structure of proteins and protein complexes in vitrified solution. Proteins tend to adsorb to the air-water interface in unsupported films of aqueous solution, which can result in partial or complete denaturation of the protein. We investigated the structure of yeast fatty acid synthase at the air-water interface by electron cryo-tomography and single-particle image processing. Around 90% of complexes adsorbed to the air-water interface are partly denatured. We show that the unfolded regions are those facing the air-water interface. Denaturation by contact with air may happen at any stage of specimen preparation. Denaturation at the air-water interface is completely avoided when the complex is plunge-frozen on a substrate of hydrophilized graphene.

## Introduction

In the short time since the resolution revolution^1^, single-particle electron cryo-microscopy (cryo-EM) has developed into a main technique for high-resolution structure determination of proteins^2^. To achieve high contrast and high resolution in cryo-EM, a small volume of protein solution is applied to an EM support grid^3, 4^ and blotted before vitrification by plunge-freezing in liquid ethane^5, 6^. This process makes it inevitable to expose the protein to the atmosphere at a high surface-to-volume ratio. It has often been suggested that the air-water interface is a hostile environment for proteins^7, 8^. Numerous studies from the first half of the 20th century^9^ have shown that globular proteins applied to dilute buffers will eventually form an insoluble monolayer of 5-10 Å thickness at the air-water interface, which means that they denature completely. A recent systematic investigation by electron cryo-tomography (cryo-ET) has shown that all of the 31 different proteins examined have a more or less pronounced tendency to adhere to the air-water interface on cryo-EM grids^10^. Even if the proteins do not denature, adsorption often results in preferential orientation, which is undesirable for image processing and high-resolution structure determination. The standard method of preparing cryo-EM specimens by plunge-freezing of thin, unsupported layers of protein solution is therefore potentially problematic. Effects of adsorption to the air-water interface on protein orientation, structure and integrity have not been investigated in detail and are not widely appreciated.

Proteins diffuse from the bulk phase of a thin layer of aqueous solution to the air-water interface in a millisecond or less^11, 12^, so that each protein can make thousands of contacts with the atmosphere during the few seconds it takes to prepare a cryo-EM grid. At each encounter, the protein is at risk of partial unfolding. Fast nanodispensers^13^ in combination with self-blotting grids^14^ have been developed to minimize protein exposure to the air-water interface, and initial results look promising^15–17.^ Attempts to overcome preferential orientation include saturation of the surface with surfactants, such as fluorinated detergents^18^ that interact poorly with the protein^19^, ^20^. These approaches require careful screening or access to a sophisticated (and costly) apparatus that is not universally available. As a simpler and more general solution, we propose to use a physical support that largely prevents protein contact with, and consequently denaturation at, the air-water interface

Continuous thin layers of amorphous carbon help to spread proteins evenly on cryo-EM grids^21–24^. Amorphous carbon is however far from ideal as a support film for cyro-EM because it adds background and conducts electrons poorly^25, 26^. Thisresults in charging and blurring of low-dose images^27, 28^. Beam-induced movement is more severe on amorphous carbon support films than on unsupported films of vitrified solutions^29^.

In contrast to amorphous carbon, graphene, a monomolecular layer of crystalline carbon, has a number of desirable properties. It is the thinnest and strongest material known and at the same time an excellent conductor^30^. It is stable under a 300 kV electron beam^31^, and almost completely electron-transparent to 2.13 Å resolution (the position of the first Bragg peak) and beyond^32^. The main problem of graphene for cryo-EM is its extreme hydrophobicity. For an even spread of the protein solution, the graphene surface has to be made hydrophilic. Graphene oxide is less hydrophobic than graphene, but also less electron-transparent^32, 33^ and it is more difficult to apply to EM grids as a monolayer^34^. Graphene can be rendered hydrophilic by plasma etching^29^ or non-covalent chemical doping, exploiting the *π*-*π* stacking interaction between graphene and aromatic planar compounds such as 1-pyrenecarboxylic acid (1-pyrCA)^35^. The advantage of non-covalent doping is that the pristine graphene surface is not modified, and that particle adsorption can be tuned by adjusting the concentration of the doping chemicals.

We used fatty acid synthase (FAS) from *Saccharomyces cerevisiae* to explore the denaturing effect of the air-water interface and how to avoid it. The structure of FAS is well-characterized by protein crystallography^36–38^ and cryo-EM^39^. This makes it easy to detect and analyze which part of the complex contacts the interface and to what extent it is denatured. We use cryo-ET to locate the FAS particles on cryo-EM grids and visualize the denaturation of individual protein complexes in contact with the air-water interface. Finally, we demonstrate by high-resolution single-particle cryo-EM that a stable substrate of hydrophilized graphene avoids the denaturation during cryo-EM specimen preparation completely.

## Results

### Fatty acid synthase is intact prior to cryo-EM grid preparation

The FAS complex used for cryo-EM data collection was pure and homogeneous, as shown by denaturing/blue-native polyacrylamide gel electrophoresis and size exclusion chromatography (Figure 1 Supplement 2 A-C). Thermal shift assays indicated that the complex was stable (Figure 1 Supplement 2 D), and at 1500-3000 mU/mg it was enzymatically fully active^40–42^. Negative-stain EM of freshly purified FAS samples indicated that the complex was structurally intact (Figure 1 Supplement 1 A). Cryo-EM of the same samples in plunge-frozen, unsupported thin layers of vitrified solution on holey carbon film revealed that around 90% of the particles had suffered major structural damage (Figure 1). FAS particles in two-dimensional (2D) and in particular 3D classification lacked between one third and one half of their density, or the density at the distal part of the beta-domes was weak (Figure 1 A, B). A reconstruction of *∼* 8,000 particles was limited to 9.5 Å resolution, according to the gold-standard 0.143 FSC criterion^43^ (Figure 1 C). Back-tracking of incomplete particles in the 3D classes (Figure 1 B) revealed major structural defects of the protein complexes in the raw micrographs (Figure 1 Supplement 1 B, C). These observations led us to conclude that the protein must have been damaged prior to or during cryo-EM grid preparation.

**Figure 1.**
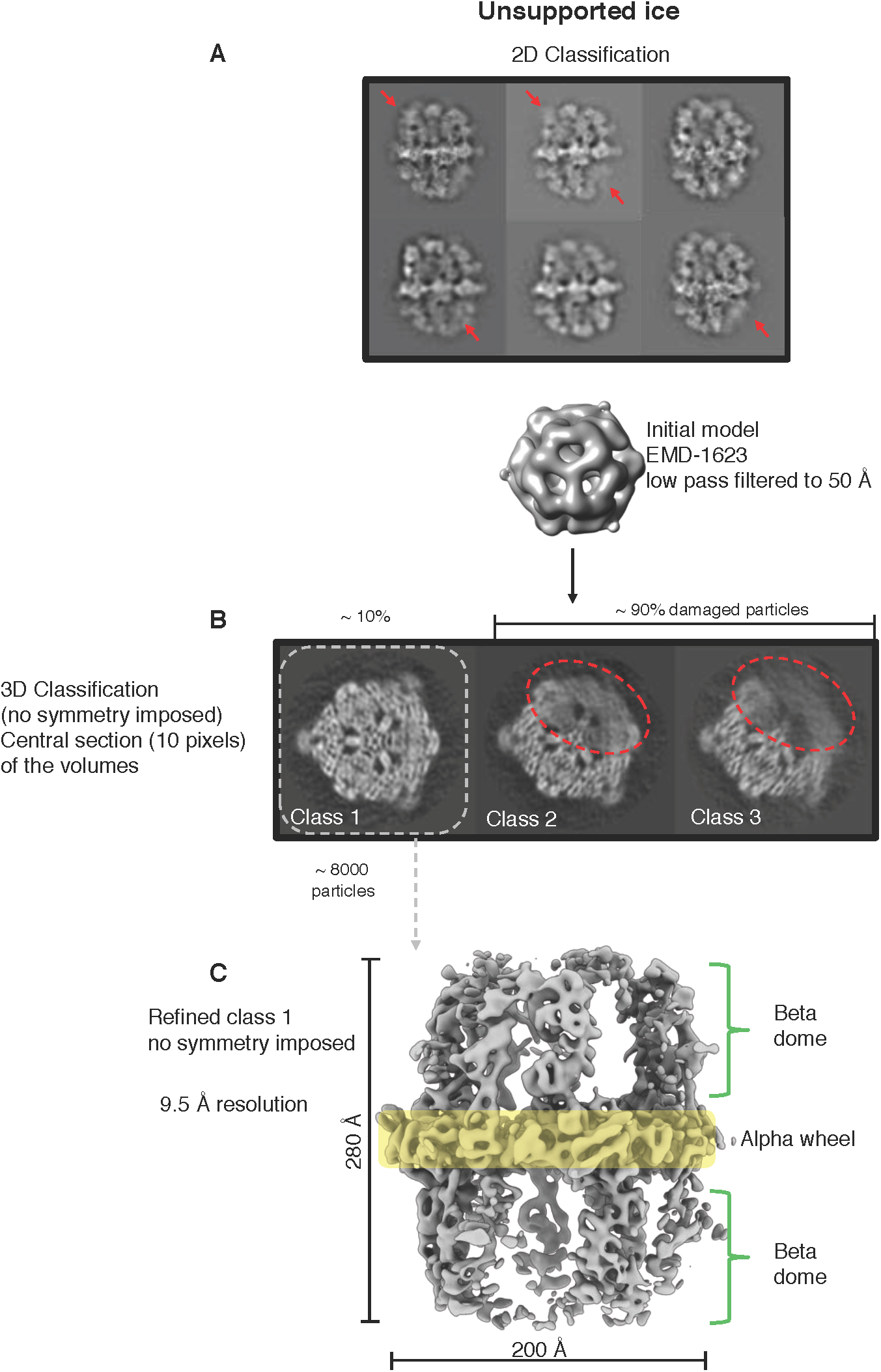
Single particle cryo-EM results from unsupported ice. (A) Two-dimensional classification of particles from unsupported vitrified solution shows class averages of FAS complexes with weak densities or absent density of beta-domes (red arrows). (B) Slices at the level of the alpha-wheel (transparent yellow in C) through the reconstructed volumes after 3D classification show major damage to about 90% of particles (dashed red). The remaining*∼*10% (dashed grey) contributed to a reconstruction (C) at 9.5 Å resolution. As starting reference of the map EMD-1623 low-pass filtered to 50 Å was used.

### Particle distribution in vitrified cryo-EM grids

Next, we performed cryo-ET on the vitrified specimens prepared for single-particle cryo-EM. Several batches of purified FAS plunge-frozen by different users under different conditions were examined. All experiments indicated damaged FAS complexes in all imaged areas (Figure 2 A). In most instances, small fragments of denatured FAS were found in the areas surrounding individual complexes (red arrows in Figure 2 A, Video S1).

**Figure 2.**
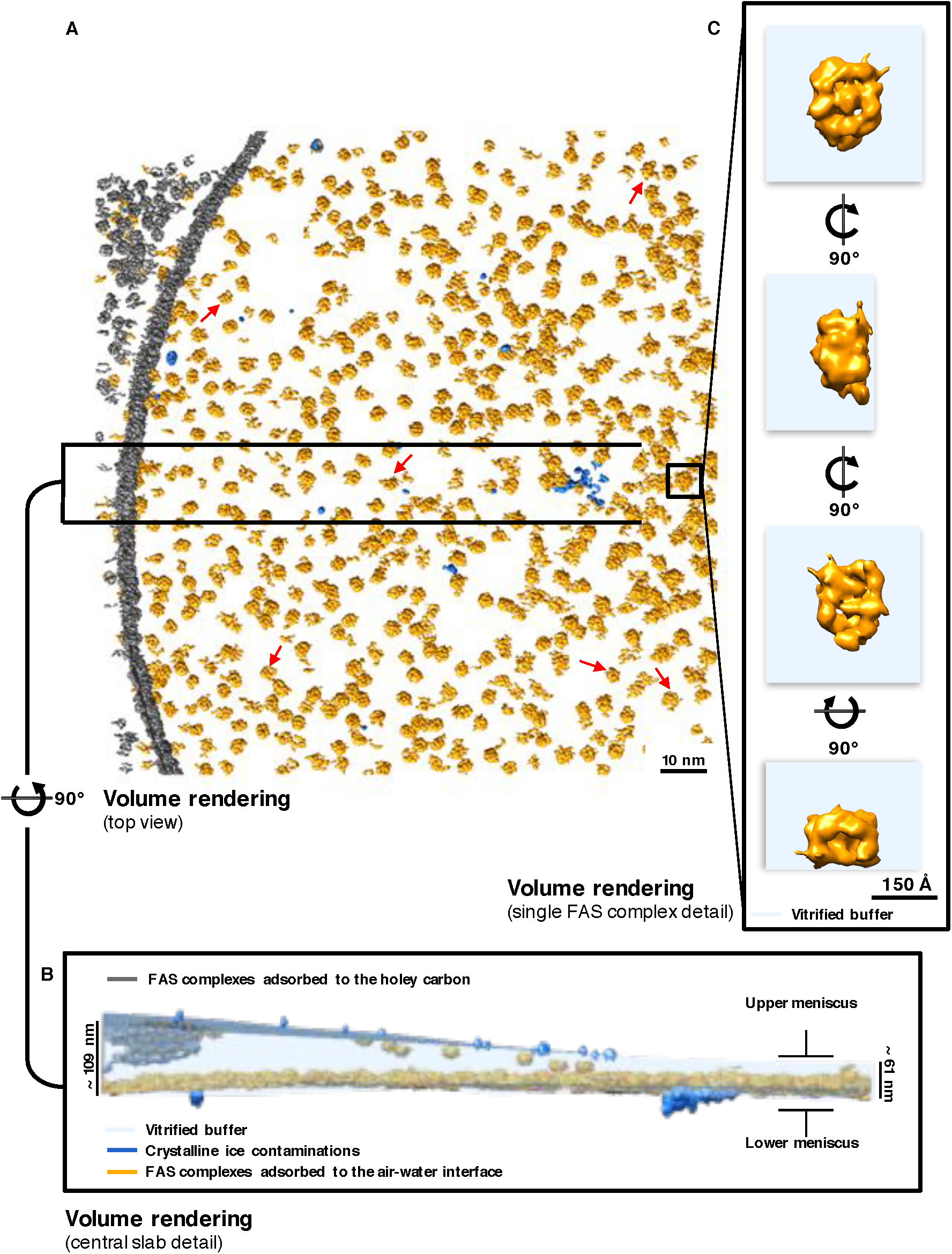
Particle distribution and structure of FAS vitrified in unsupported vitrified solution. ((A) Overview of a typical Quantifoil R2/2 grid hole, containing FAS complexes vitrified without support, visualized by convolutional neural network segmentation. Red arrows indicate FAS fragments. (B) Slab of vitrified buffer, delimited by carbon and contaminating atmospheric ice crystals. (C) Detail of a single FAS molecule showing morphological difference between the sides facing the vitreous buffer or the air-water interface.

Cryo-ET revealed that FAS adhered to the two opposite surfaces of the unsupported thin layers of vitrified solution. One surface, which we refer to as the lower meniscus, was densely packed with adsorbed protein complexes. The opposite surface (the upper meniscus) had only a small number of particles attached (Figure 2 B and Figure 2 Supplement 1). Together with small ice crystals from atmospheric contamination on the outside surface of the vitrified layer, the FAS complexes on the upper and lower meniscus allowed us to trace the air-water interface exactly (Figure 2 B).

Tomographic volumes indicated that most of the FAS particles in contact with the air-water interface were damaged. The particles were mostly flattened on one side and appeared incomplete (Figure 2 C). The flattened regions of these partly denatured particles aligned with the plane of the air-water interface. Particles attached to the lower and upper meniscus appeared to be equally affected, although the small number of particles on the upper meniscus precluded a statistically significant analysis. Our observations thus suggest that at some point during cryo-specimen preparation, the large majority of FAS complexes encountered the air-water interface, attached to it, and the air-exposed side unfolded before vitrification.

### Orientation of damaged FAS particles on the air-water interface

The extent of structural damage to the FAS particles at the air-water interface was investigated by subtomogram averaging (STA). A set of 1,724 subvolumes was manually selected, and a subset of 20 randomly picked volumes was used as a reference for initial alignment. No symmetry constraints were applied. The final reconstruction indicated that one side of the FAS map lacked density, whereas the opposite side of the complex appeared intact (Figure 3 A). FAS attached always with its long axis parallel to the air-water interface, which accounts for the scarcity of top views in the single-particle analysis. To determine the orientation of the partly denatured FAS complexes relative to the air-water interface, we fitted a surface through the centers of all particles (Figure 3 B). We then calculated the vectors pointing from the center of a complex towards its flattened side (Figure 3 C). Finally, we assessed by how much the vectors diverged from the normal of the previously calculated plane through all particles at that position, and whether they pointed towards the air-water interface or away from it. This analysis indicated clearly that the vectors pointed towards the air-water interface (Figure 3 D) (see Materials and Methods).

**Figure 3.**
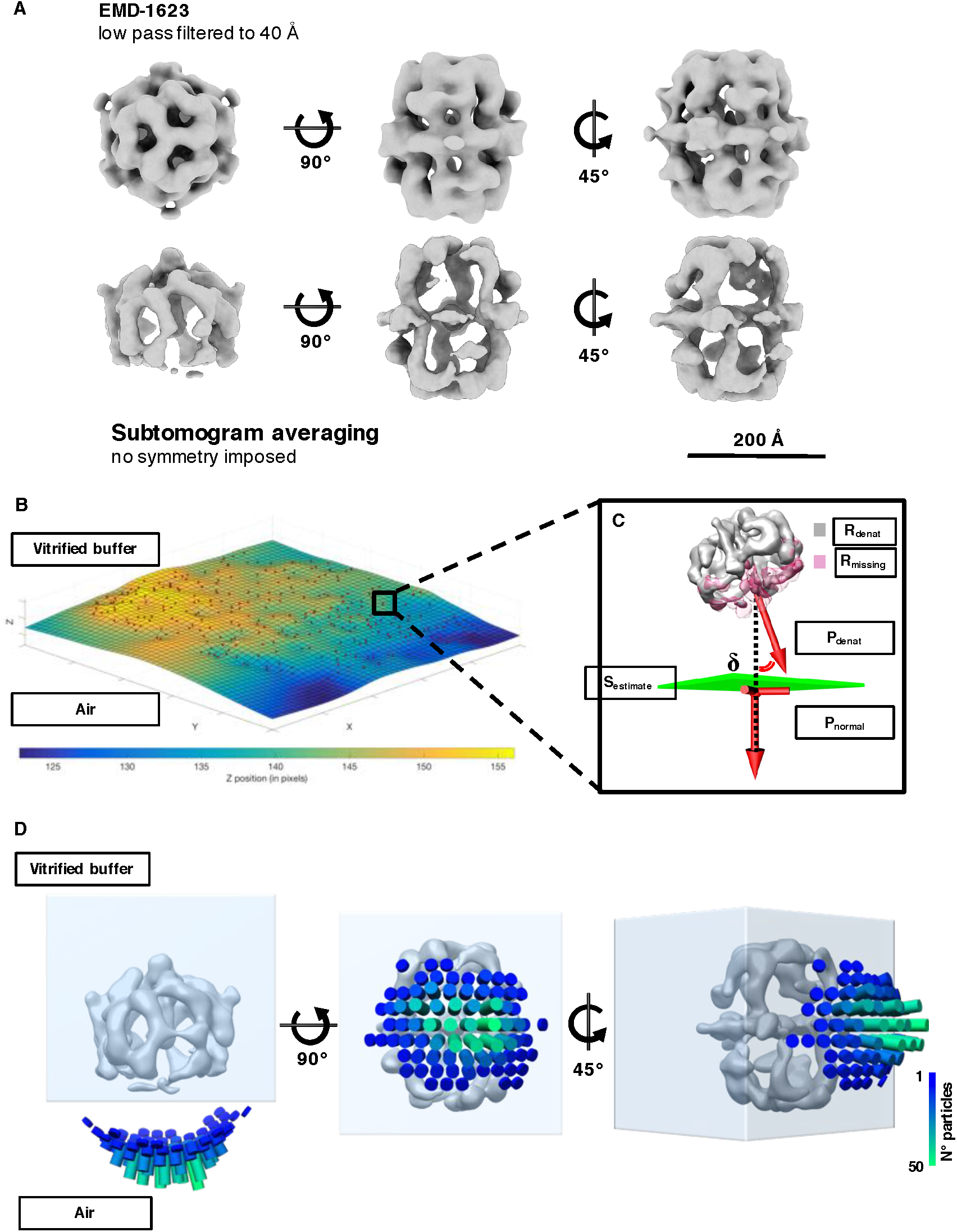
Sub-tomogram averaging and orientation of denatured FAS in unsupported vitrified solution. (A) Subtomogram averaging of FAS complexes in unsupported vitrified solution. Denaturation of the complex is apparent on one side, while the opposite side is undamaged. A map of intact (EMD-1623) is shown above for comparison. (B) Plot of a thin plate spline function representing the surface passing through the coordinates of the centers of the particles (S_estimate_). The orientation of the particles with respect to the interface was determined with the displacement angle *δ* (C) between the vector (P_denat_) pointing from the center of the broken particle (R_denat_) to the center of mass of the missing part (R_missing_), and the vector normal to the surface (P_normal_) in the same coordinate point. (D) Total distribution of *δ* from all the particles adsorbed to the most populated ice meniscus of all the tomograms.

The structural heterogeneity of the partly denatured FAS complexes was examined by multi-reference alignment (see Materials and Methods). In line with the single-particle results (Figure 1 B), we found different degrees of particle damage. About 86% were extensively damaged, with one third or even half of the characteristic quaternary FAS structure weak or absent (Figure 3 Supplement 1 A). The remaining 14% had poorly resolved densities (Figure 3 Supplement 1 B), suggesting that even those particles on the air-water interface that appeared intact had suffered some damage. The subvolume set probably contained a small number of undamaged particles from the bulk phase, but visual inspection of the tomographic volumes did not reveal any. We conclude that most if not all particles at or near a meniscus were damaged to a greater or lesser extent by contact with the air-water interface.

### Air exposure induces protein denaturation

In a series of three experiments we tested different ways in which exposure to air could cause protein denaturation (see Materials and Methods). As before, negative-stain EM confirmed that the particles were initially undamaged (Figure 4 A). In one experiment we bubbled air through the sample to maximize air contact. In another experiment we poured the protein solution over a glass rod^8^ to expose a continuous thin film to the atmosphere (Figure 4 B). In the third experiment we applied a 20 *µ*l volume of FAS solution to a standard EM support grid coated with continuous carbon and touched the top of the droplet with a second carbon-coated grid (Figure 4 C). In this way we separated the particles adsorbed to the air-water interface from those adsorbed to the carbon film (Figure 4 D). The result of each experiment was then examined by negative-stain EM (upper panels in Figure 4 B to D). Bubbling air through the sample (experiment 1) completely denatured all FAS complexes (not shown), whereas in experiments 2 and 3 a small proportion remained intact. Denatured proteins were a predominant feature in all the three conditions except that particles adsorbed to the carbon film in experiment 3 (Figure 4 D) were apparently undamaged. These results show that FAS at the air-water interface is denatured, whereas it remains intact when adsorbed to a solid substrate in liquid.

**Figure 4.**
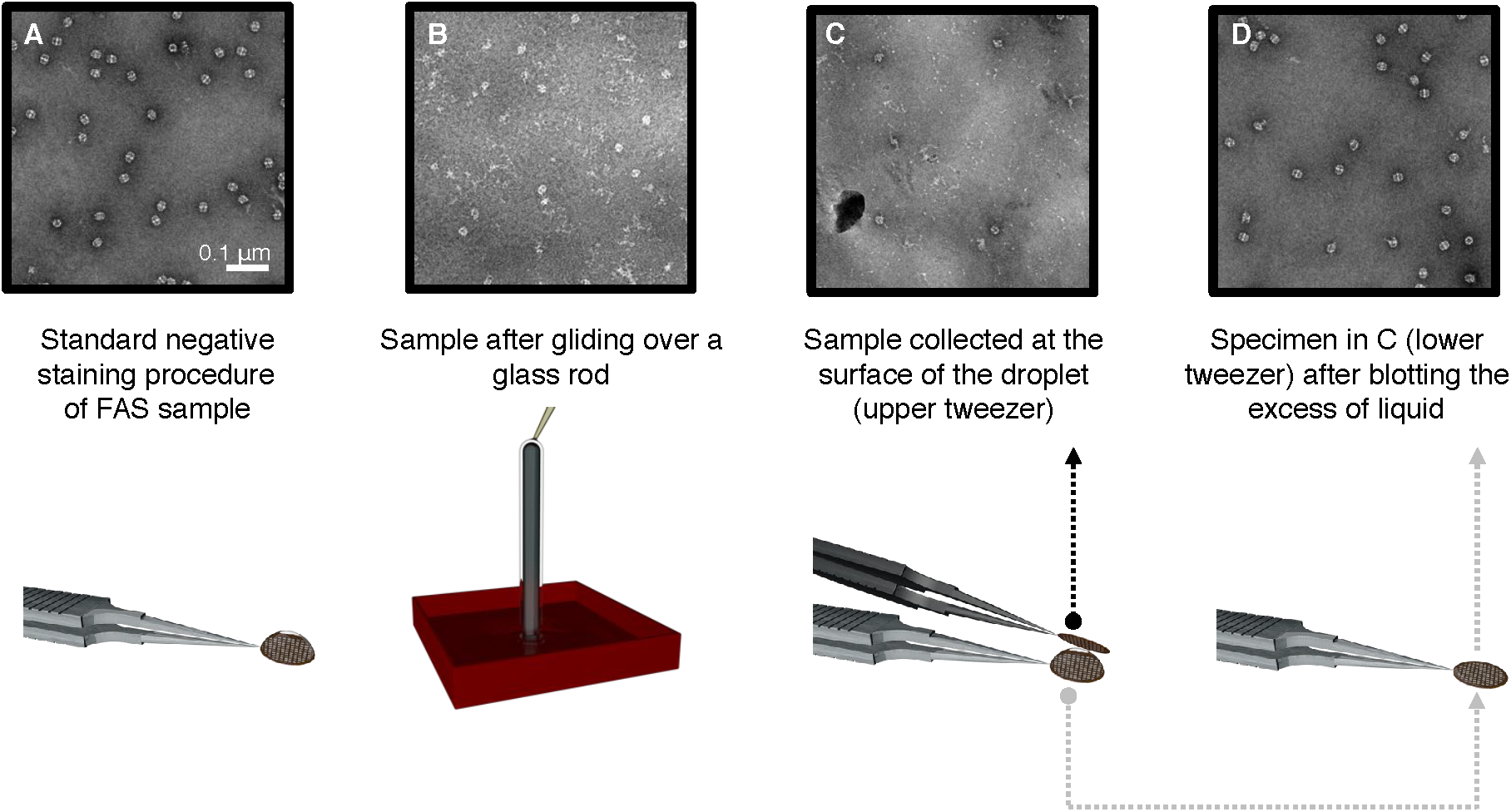
Denaturation by controlled air exposure. (A) Negative stain control of untreated FAS. (B) A thin film of FAS solution flowing over a glass rod. Negative staining reveals that most complexes are denatured. (C) A drop of FAS solution was placed on a carbon-coated EM grid (lower tweezers) and touched with another carbon-coated grid (upper tweezers) to pick up protein complexes at the drop surface. (D) Negative staining of the lower grid in C indicates only intact FAS complexes. Three-dimensional rendering with Blender.

### Hydrophilized graphene-coated grids prevent denaturation at the air-water interface

To find out whether adsorption to a continuous support film would prevent damage also under cryo-conditions, we prepared FAS on EM-grids coated with a layer of graphene rendered hydrophilic with 1-pyrene-carboxylic acid (1-pyrCA). To assess the quality of the graphene, all grids were examined by electron diffraction before vitrification. Sharp diffraction spots indicated flat monolayers of graphene (Figure 5 A, B). The hydrophobic nature of the untreated graphene film was apparent from the repulsion of a water droplet pipetted onto the grid (Figure 5 C). The same grids were then chemically doped with a solution of 1-pyrCA, which did not degrade the crystalline order of the graphene layer (Figure 5 D, E). The hydrophilic character of the 1-pyrCA-doped graphene was indicated by the reduced contact angle of a water droplet on the grid (Figure 5 F). The FAS solution was applied as before, and grids were blotted and plunge-frozen as for unsupported vitreous films. The hydrophilized graphene/FAS grids were then used for cryo-ET and single-particle cryo-EM.

**Figure 5.**
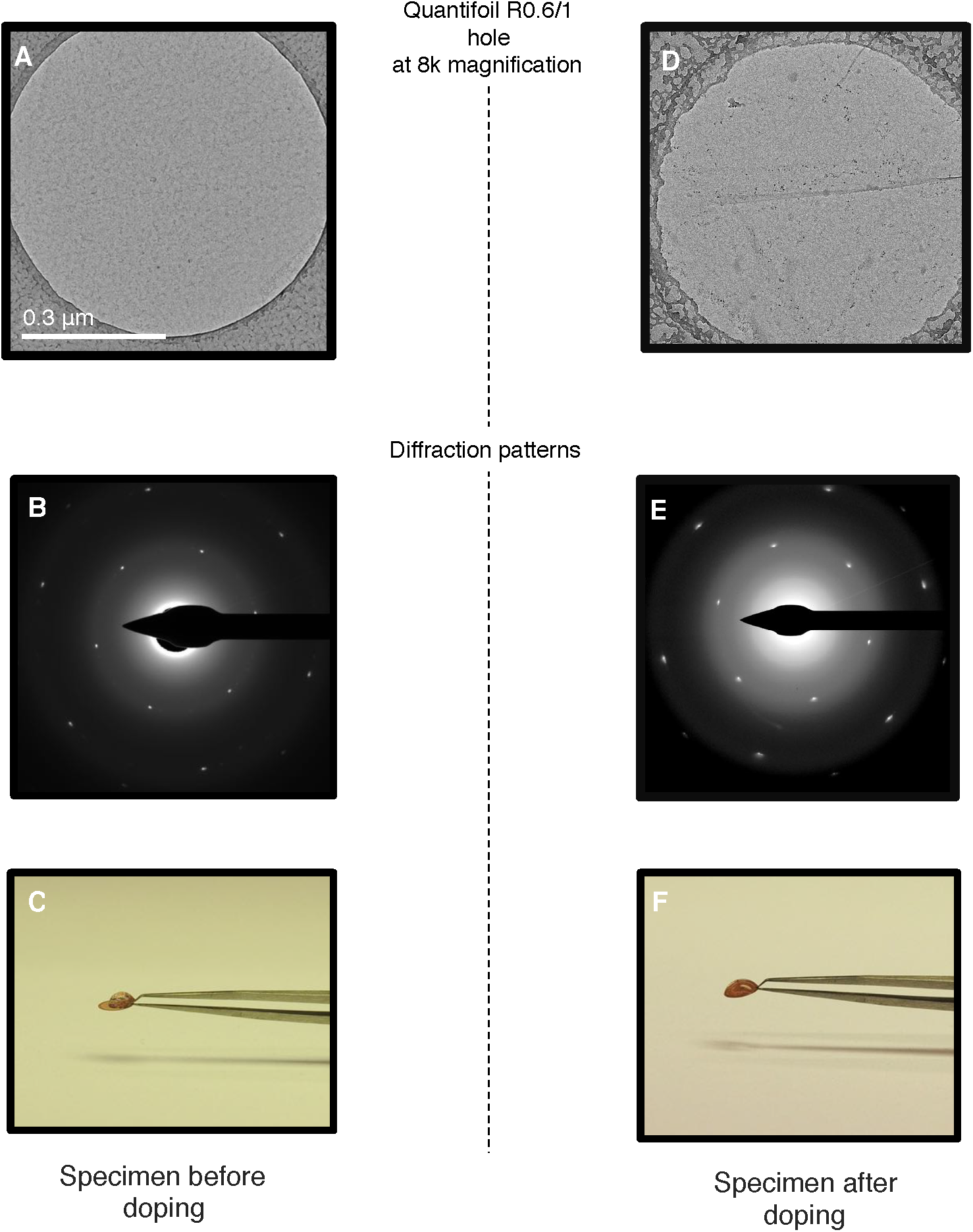
Chemical doping of graphene-coated Quantifoil grids. (A) Overview of a hole from a Quantifoil R0.6/1 grid coated with graphene. (B) Electron diffraction pattern of the area shown in (A). (C) Same grid with a droplet of water. (D) Same grid as in (A) after chemical doping with 1-pyrCA. (E) Electron diffraction pattern of the area imaged in (D). (F) 1-pyrCA-doped grid with a droplet of water.

Cryo-ET indicated the position of the air-water and graphene-water interfaces by atmospheric ice crystals and small patches of contaminants (Figure 6 A, B). When the graphene layer was rendered hydrophilic by 1-pyrCA, FAS had a strong preference for the graphene-water interface (Figure 6 B, Figure 6 Supplement 1), and only very few particles were detected at the air-water interface (Video S2).

**Figure 6.**
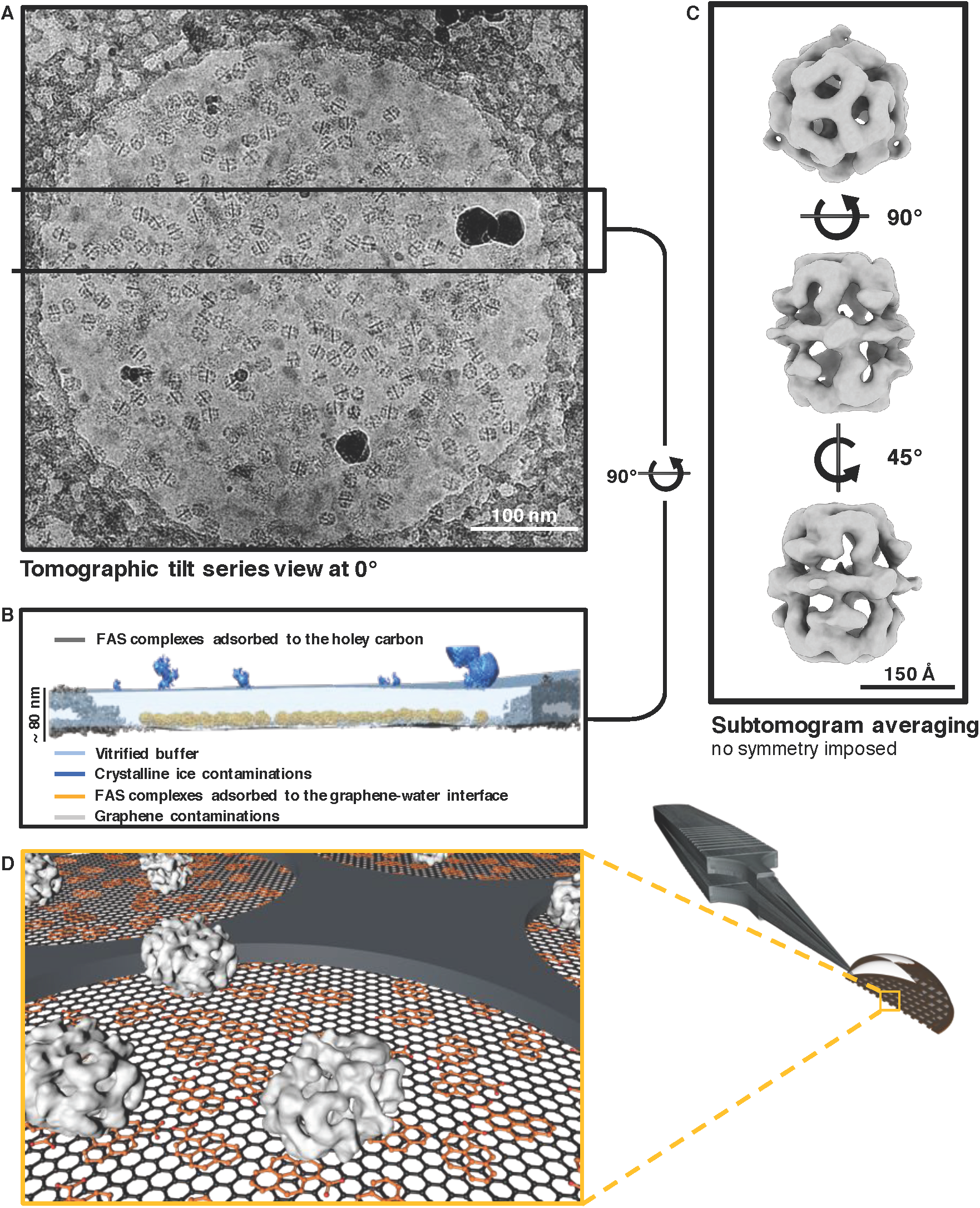
STA of FAS vitrified on hydrophilized graphene. (A) Low pass-filtered (10 Å frequency) micrograph of a typical hole covered with hydrophilized graphene and FAS applied. (B) Slab of vitrified buffer on hydrophilized graphene with FAS complexes below and atmospheric ice contamination on top, delineating the upper meniscus. (C) Subtomogram averages of FAS complexes vitrified on hydrophilized graphene. (D) Schematic graphical representation (Blender) of grid holes: Quantifoil holey carbon (dark grey), graphene monolayer (black hexagonal mesh), 1-pyrCA (dark orange), FAS complexes (light grey). Individual components are not drawn to scale.

To investigate the state of preservation of FAS on hydrophilized graphene, we hand-picked a set of 2,090 subvolumes and performed subtomogram averaging and multi-reference classification as for unsupported vitrified samples. Reconstructions both before (Figure 6 C) and after (Figure 6 Supplement 2) multi-reference alignment indicated that all particles were intact. The best sub-tomogram averages yielded maps at 24.6 Å and 17.1 Å resolution before and after masking (Figure 6 Supplement 3). Since few if any particles stuck to the air-water interface and multi-reference alignment did not reveal any damage, we conclude that FAS does not denature on hydrophilized graphene.

### Hydrophilized graphene is suitable for high-resolution cryo-EM

To find out whether hydrophilized graphene films are suitable for high-resolution structure determination, we analyzed FAS on these grids by single-particle cryo-EM. Typical micrographs recorded at 0.9 *µ*m defocus showed good contrast (Figure 7 A). The corresponding 2D (Figure 7 B) and rotationally averaged 1D power spectra (Figure 7 C) generated from the particles only (since graphene is electron-transparent in this resolution range) indicated oscillations beyond 3 Å spatial frequency.

**Figure 7.**
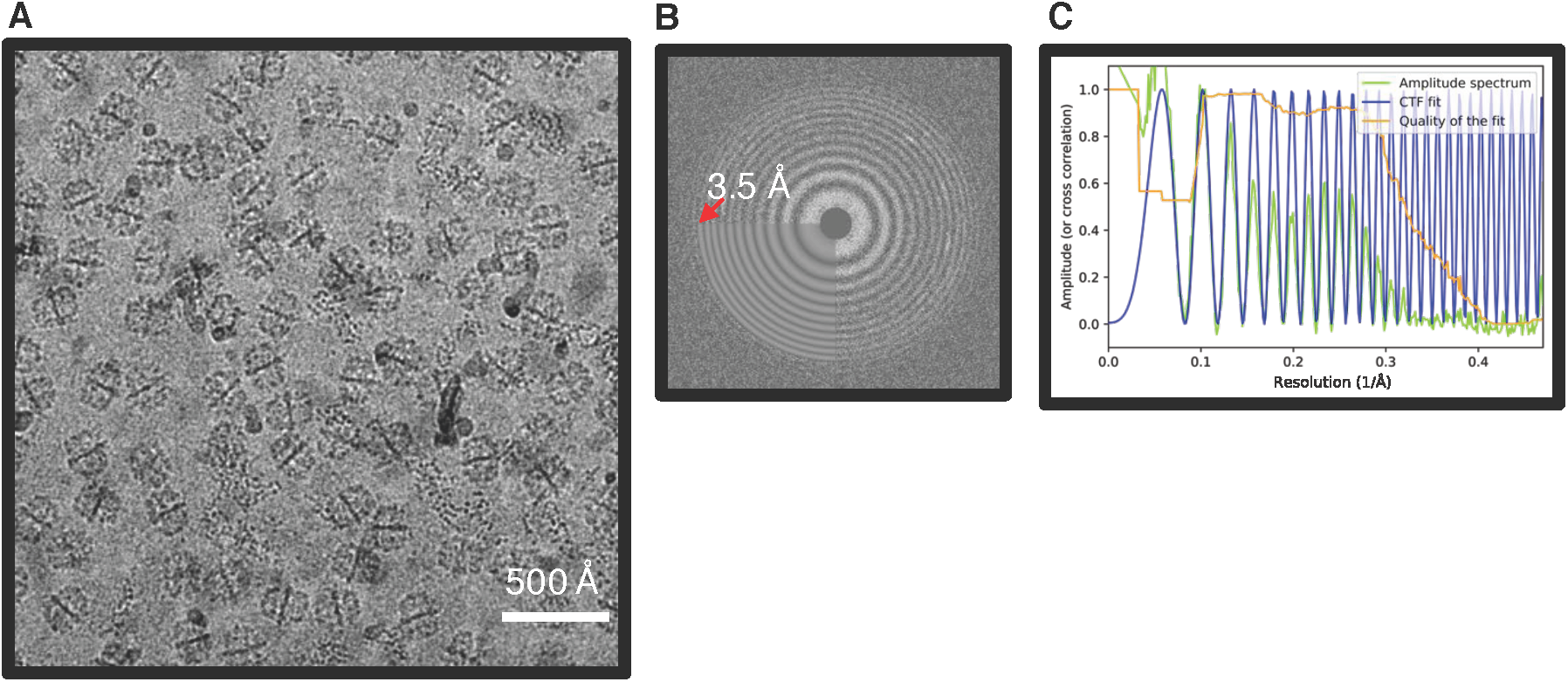
Quality of the graphene hydrophilized grids in cryo-conditions. (A) Typical area of FAS particles on Quantifoil R0.6/1 grid coated with a monolayer of graphene imaged at 0.9 *µ*m defocus low-pass filtered to 5 Å. (B) Oscillations of the corresponding experimental 2D power spectrum extend beyond the highest frequency used to fit the theoretical CTF (3.5 Å). (C) A one-dimensional plot of the power spectrum shows that the CTF fit extends beyond a spatial frequency of 3 Å.

All 2D class averages displayed high-resolution detail (Figure 8 A) and confirmed that FAS was structurally undamaged. This was confirmed by 3D classification, which showed intact complexes with well-resolved secondary structure. Note that this dataset contained only intact particles (Figure 8 B), whereas 90% of the particles in the single-particle FAS dataset from unsupported vitrified samples had suffered major damage (Figure 1 B classes 2,3), and even the remaining 10% were compromised (Figure 1 B class 1 and Figure 1 C).

**Figure 8.**
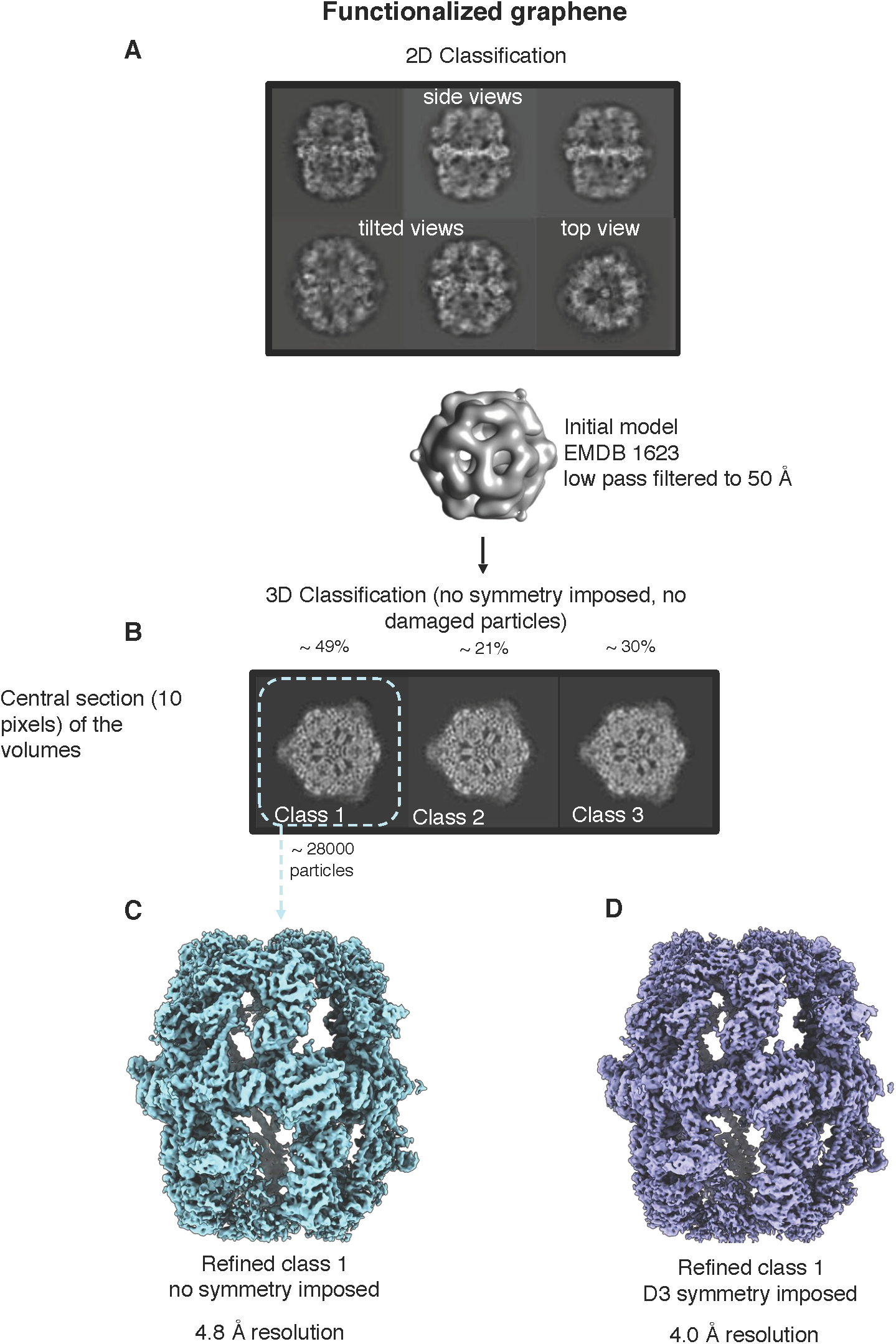
Single-particle cryo-EM of FAS on hydrophilized graphene. (A) Two-dimensional class averages of FAS on hydrophilized graphene show intact particles. The 3D map of FAS in unsupported vitrified buffer (EMD-1623) was used as an initial reference. (B) 3D classification revealed that all particles are intact, in agreement with STA results (Figure 6 Supplement 2 A). A final map calculated without (C) or with imposed D3 symmetry (D) indicated a resolution of 4.8 Å or 4.0 Å.

After merging the best 3D classes, we obtained a final set of *∼*28,000 particles. Auto-refinement without symmetry (C1) or with imposed D3 symmetry yielded maps at 4.8 and 4.0 Å resolution, respectively (Figure 8 C, D). Local resolution estimates indicated better than 3.5 Å resolution for the rigid alpha wheel (Figure 8 Supplement 1). For an unbiased comparison to the 9.5 Å map obtained from FAS in unsupported vitrified buffer (Figure 8 Supplement 2 A), we randomly selected 8,000 particles from the dataset collected on hydrophilized graphene. The resulting map attained a resolution of 6.4 Å (Figure 8 Supplement 2 B), confirming that hydrophilized graphene works very much better as a substrate for single-particle cryo-EM of FAS than unsupported vitrified buffer. The particles were intact and the map contained all the main features of the best non-symmetrized 4.8 Å map (Figure 8 Supplement 2 C).

## Discussion

An earlier single-particle cryo-EM structure of *S. cerevisiae* FAS in unsupported vitrified samples^39^ reported a resolution of 7.2 Å at the 0.5 FSC threshold and 5.9 Å at 0.143 FSC. The “gold-standard FSC” procedure of estimating map resolution by comparing reconstructions derived from two independent halves of the particle data set^43, 44^ had not been introduced at the time and was not applied. Therefore, the 0.5 FSC resolution estimate was more realistic, as confirmed by a comparison of the FAS alpha wheel in the earlier structure to the gold-standard FSC maps in the present study (Figure 8 Supplement 3). The resolution of the earlier map is clearly between that of the 6.4 Å map of FAS on graphene and the 9.5 Å map of FAS in unsupported vitrified buffer, which were obtained with *∼*8,000 particles each. The better quality of the earlier map^39^ is fully accounted for by the larger number of particles contributing to it, and the application of D3 symmetry. The fact that the resolution was limited to 7.2 Å even with more than twice the number of particle images suggests that the complex was equally affected by denaturation at the air-water interface. The 4 Å resolution of our present map from 28,000 particles on hydrophilized graphene is most likely limited by the inherent flexibility of the complex. This is implied by the cryo-EM structure of a FAS complex from the thermophilic fungus *Chaetomium thermophilum*^45^, which attained a gold standard, FSC 0.143 resolution of 4.7 Å from only ∼4,000 particles. It is well known that protein complexes from thermophilic organisms are more stable than those of mesophilic origin. By comparison, more than 100,000 particles contributed to the 7.5 Å cryo-EM structure of a FAS complex from *Mycobacterium smegmatis*^46^, suggesting that the mycobacterial complex is significantly less stable.

In many of the cryo-EM grids examined by electron tomography^10^, the curvatures of the upper and lower meniscus were different. In the case of FAS, the less densely populated upper meniscus was more strongly curved than the lower meniscus (Figure 2 B, Video S1). The reason for the difference in curvature is not known and most likely stochastic. The asymmetric distribution of protein on the upper and lower meniscus is surprising, because the grids were blotted symmetrically from both sides. Possibly the more densely populated lower meniscus remained in contact with air through the blotting process, so that more protein accumulated on it. Alternatively, protein that would have adhered to the upper meniscus may have been scavenged by adsorption to the glow-discharged holey carbon film. At this stage, the reason for the asymmetrical particle distribution on the two surfaces is unknown.

Our cryo-EM analysis of FAS, a soluble 2.6 MDa protein complex, revealed that only a minority of the particles in unsupported vitreous films retained intermediate-resolution features, whereas up to 90% were at least partly denatured by the air-water interface. A survey of recently reported high-resolution cryo-EM structures shows that usually only a minor fraction of a large single-particle data set contributes to the final high-resolution map. Percentages of good particles were 19% for the3.8 Å map of a human synaptic GABAA receptor^47^; 15% for human P-glycoprotein at 3.4 Å^48^; 11.8% for a 4 Å nucleosome map^49^; 8.9% for the 3.4 Å structure of human γ-secretase^50^; and only 5.7% for the 4 Å structure of a sodium channel complex from electic eel^51^. All were prepared in unsupported vitrified buffer. These numbers suggest that up to 94% of the particles may have suffered partial denaturation at the air-water interface. Future studies will show whether this is indeed the case, and whether denaturation can be avoided by using hydrophilized graphene grids, as we have shown for yeast FAS. If hydrophilized or otherwise functionalized graphene grids prove to work as well for other proteins to overcome denaturation at the air-water interface, a much higher proportion of particles would contribute to the final structure. This would result in a large increase in data collection efficiency and significantly better maps. It would be a major boost for cryo-EM.

## Methods

- **Strain cultivation and protein purification** Yeast cultivation and FAS purification were carried as previously reported^52, 53^. Briefly, haploid FAS-deficient S. cerevisiae strain BY.PK1238_KO cells were transfected with plasmids carrying the FAS-encoding genes pMF319d(pRS313_FAS2) and pMF639 (pRS315_FAS1-twinStrep-II), then grown in YPD medium until the optical density OD600 reached *∼*11. After cell harvesting and disruption by beating glass beads, the insoluble components were eliminated by centrifugation and the supernatant was purified through strep-Tactin affinity chromatography. The strep-tagged FAS was eluted and concentrated with a 100 kDa cutoff filter, then further purified by size-exclusion chromatography. The main peak was finally concentrated to *∼* 4 mg/mL. All purification steps were analyzed by SDS-PAGE.
- **Thermal shift assay (TSA) and activity assay** Thermal shift assays were performed as previously reported^54^. Briefly, 2 *µ*L of protein solution (0.9 mg/mL) were mixed with 21 *µ*L of phosphate buffer (100 mM; pH 6.5) and 2 *µ*L of 62.5 X SYPRO Orange protein gel stain, then fluorescence was measured from 5 °C to 95 °C with a step of 0.5°C/min, with excitation wavelength set to 450-490 nm, and emission wavelength to 560-580 nm. FAS activity was determined by tracing NADPH consumption at 334 nm as previously reported by Gajewski et al., 2017^55^, except that it was adapted for platereader read-out (120 *µ*L scale containing 200 mM NaH_2_PO_4_/Na_2_HPO_4_ (pH 7.3), 1.75 mM 1,4 dithiothreitol, 0.03 mg/mL BSA, 0.7 *µ*g FAS, 500 *µ*M malonyl CoA, 417 *µ*M acetyl CoA and 250 *µ*M NADPH).
- **Grid preparation** Quantifoil 0.6/1 and R2/2 grids (Quantifoil Micro Tools, Jena, Germany) were used to prepare cryo-specimens with or without graphene support. Grids were washed thoroughly overnight in chloroform before use. For graphene-coated grids, graphene pads (1 cm^2^) (Graphenea, Cambridge, MA) which were floated onto Quantifoil grids in a water bath. Quantifoil R1/2 and R1.2/1.3 grids were also tested but the smaller hole size yields flatter graphene layers. Grids were air-dried for 30 minutes and then heated to 150°C for one hour to anneal the graphene layer to the Quantifoil film. Graphene-coated grids were stored under vacuum over night. The grids were washed in pure acetone and isopropanol for one hour each and dried under a nitrogen stream. Finally, graphene-coated grids were dipped into 50 mM 1-pyrenecarboxyilic acid (Sigma Aldrich, Munich, Germany) dissolved in DMSO (Sigma Aldrich, Munich, Germany). Grid quality was assessed in a FEI Tecnai G2 Spirit BioTwin (FEI Company, Hillsboro, OR) operated at 120 kV, at a nominal magnification of 9,300x, yielding a pixel size at the specimen level of 1.19 nm. Electron diffraction patterns were recorded at a nominal camera length of 540 mm, with a 1 s exposure time and 150 *µ*m aperture.
- **Negative Stain** FAS was diluted in purification buffer (100 mM sodium phosphate, pH 6.5) to a final concentration of 0.05 mg/ml and negatively stained with 2% (w/v) sodium silicotungstate (Agar Scientific, Stansted, UK). Three *µ*l of protein solution was applied onto freshly glow-discharged carbon-coated copper grids. After blotting the excess of protein solution, the staining solution was immediately applied to the grid and blotted off from the same side. This step was repeated twice. Micrographs were recorded in a FEI Tecnai G2 Spirit (FEI Company, Hillsboro, OR) operated at 120 kV, at a nominal magnification of 42,000x, yielding a pixel size at the specimen level of 2.68 Å.
- **Controlled protein denaturation at the air-water interface** Triplicates of three different experiments of controlled protein denaturation at the air-water interface were carried out. Freshly purified FAS solution was diluted to 0.01 mg/ml with purification buffer (100 mM sodium phosphate, pH 6.5). For experiment 1, air was bubbled through 200 *µ*L of protein solution through a pipette tip for about 10 seconds, and EM grids were prepared from a 3 *µ*L aliquot. For experiment 2, 200 *µ*L of protein solution were passed over a 5 cm 100 *µ*L intraMARK disposable glass micropipette (Brand, Wertheim, Germany) sealed at both ends, and collected in a 1.5 ml tube for EM analysis in negative stain. For experiment 3, 20 *µ*L of FAS solution were pipetted onto EM grid coated with amorphous carbon and incubated in air. After 15 seconds, the drop was touched with a second carbon-coated EM grid, blotted and negatively stained as before.
- **Single particle cryo-EM** Three *µ*l of FAS solution (concentration 1.0 mg/ml or 0.2 mg/ml for unsupported grids or hydrophilized graphene grids respectively) were applied to freshly glow-discharged Quantifoil R2/2 holey carbon grids (Quantifoil Micro Tools, Jena, Germany) or Quantifoil R0.6/1 holey carbon grids for hydrophilized graphene. The grids were vitrified in a Vitrobot Mark IV plunge-freezer at 100% humidity and 10°C after blotting for 6-8 s. Cryo-EM images were collected in a Titan Krios (FEI Company, Hillsboro, OR) electron microscope operating at 300 kV. Images were recorded automatically using the software EPU at a nominal magnification of 135,000x, yielding a calibrated pixel-size of 1.053 Å, on a Falcon III EC direct electron detector (FEI Company, Hillsboro, OR) operating in counting mode. Dose-fractionated 90 s movies of 81 frames were recorded at a dose rate of 0.4 e^-^/pixel/s (*∼* 32 e^-^/Å^2^ total dose at the specimen) and a defocus range between 0.8-2.5 *µ*m. A total of 2648 movies were collected of FAS in unsupported vitrified buffer, and 1055 movies of FAS on hydrophilized graphene. Whole-image drift correction and magnification distortion correction of each movie was performed using Unblur^56^. A second round of local drift-correction was performed using MotionCor2^57^, dividing each frame in 5×5 patches and applying a dose-dependent weighting with the scheme implemented by Grant and Grigorieff, 2015^56^. The CTF was determined on the aligned movies using CTFFIND 4.1.10^58^ averaging 10 frames at a time in order to reach *∼*4 e^-^/Å^2^electron dose^59^. Unless specified otherwise, all subsequent image processing steps were performed within Relion 2.1^60^. For unsupported vitrified solutions, a set of 600 manually picked particle images (box size 364 x 362 pixels) was subjected to 2D reference-free classification to generate a first reference for automatic particle picking. Automatic picking resulted in an initial dataset of 82,353 particles, which was 2D classified to remove false positive picks. The remaining 81,163 particles contributed to the reconstruction of the first consensus refinement using a FAS EM map (EMD-1623^39^) low-pass filtered to 50 Å as initial reference without imposing symmetry. The same procedure was applied to the data collected on hydrophilized graphene grids, resulting in a set of 57,467 and 57,021 particles after autopicking and 2D classification respectively. Extensive 3D classification with local angular search using 3 to 9 classes was applied to select the particles that contributed to the best 3D classes. The best 28,132 particles, sorted by 3D classification, were combined to perform a reconstruction imposing C1 (no symmetry) or D3 symmetry. This yielded maps at 9.5 Å resolution (unsupported vitrified solution, C1 symmetry), 4.8 and 4.0 Å resolution (hydrophilized graphene, C1 and D3 symmetry, respectively). All maps were corrected for the modulation transfer function (MTF) of the Falcon direct detector and sharpened by applying a B-factor of −130 Å^2^. Local resolution was estimated using Relion 2.1^60^. All the 3D rendering were performed with ChimeraX^61^.
- **Electron cryo-tomography** Vitrified specimens were imaged with a Titan Krios (FEI Company, Hillsboro, OR) electron microscope operating at 300 kV, equipped with a K2 summit direct electron detector and a Quantum energy filter (Gatan, Inc., Pleasanton, CA). The nominal magnification was set to 42,000x or 64,000x, yielding a calibrated pixel size at the specimen height of 3.39 Å or 2.20 Å for the dataset in unsupported aqueous films or hydrophilized graphene respectively. The illumination was set to a fluence of *∼*6-9 e^-^/pixel/s. Tomographic images were automatically recorded in counting mode using a dose-symmetric acquisition scheme^62^ implemented in SerialEM^63^, with a defocus of −5 to −7 *µ*m.Dose-fractionated images were recorded from −60°to +60° with a tilt step of 3°, and a cumulative dose of 90 e^-^/Å^2^ per tilt series. After movie frame alignment with MotionCor2^57^, the images were CTF corrected with the software Gctf^64^. Stack files generated from single micrographs were selected as input for IMOD^65^, where all views were aligned automatically through patch tracking and used to reconstruct 3D volumes with a weighted back-projection algorithm. If necessary, the reconstructed volumes were processed with a Nonlinear Anisotropic Diffusion (NAD) filter^66^ for visualization.
- **Subtomogram averaging, volume segmentation and rendering** All processing and analyses of 3D particles for structure determination were performed with Dynamo^67^. A set of 1724 and 2090 particles was manually picked from unsupported and hydrophilized graphene-coated grids respectively, avoiding the ones adsorbed to the surface of the holey carbon. In both cases a subset of 20 random particles was used to generate an initial reference, then the signal-to-noise ratio increased through subtomogram averaging. For both dataset the particles were split into two independent subsets, which were aligned to two references. Finally, the heterogeneity was further explored by means of multi-reference alignment, where clones of the same map are convoluted with Gaussian noise of equal amplitude and provided as initial reference for the classification. In order to exclude reference bias during this procedure we provided initial models bearing the opposite structural features of analyzed particle dataset: the STA resulting from unsupported aqueous layers (damaged FAS complex) was used for our dataset on graphene, and the same map used for single particle (EMD-1623, intact FAS complex) for the one from unsupported samples. Both references were low-pass filtered to 50 Å prior to convolution with noise. The final map was band-pass filtered to 308 and 12 Å frequencies. For gold-standard resolution estimation, 0.143 FSC value was used. To assess mask bias, FSC was also performed on the masked half-maps with phases randomized beyond 60 Å. The correlation drops at the resolution above which the phases were randomized, indicating that the mask does not affect the resolution estimate (Figure 6 Supplement 3). For illustrative purposes the final maps were Gaussian-filtered (standard deviation of two physical pixels) within UCSF Chimera^68^ and the tomographic volumes segmented with the convolutional neural network method implemented by Chen et al. 2017^69^ within the EMAN2.2 software package^70^. Graphical rendering and movie editing were performed with UCSF Chimera^68^ and ChimeraX^61^.
- **Estimation of particle-to-interface orientation** To determine the orientation of partly denatured FAS with respect to the air-water interface, MATLAB was used to correlate the tomographic reconstruction with a geometrical model assuming that, upon adsorption, the plane describing the denatured side of the FAS particles would be parallel to the air-water interface. The missing density due to denaturation (R_missing_) was treated as the difference between the reconstruction of intact FAS (R_intact_) and the map obtained from sub-tomogram averaging of denatured particles (R_denat_). The analysis consisted of five sequential steps: (i) coordinates of the center of FAS complexes previously determined via sub tomogram averaging were used to model the air-water interface (S_estimate_) using the *Thin-plate interpolator* option of the *Curve Fitting* Toolbox in MATLAB (Figure 3 C); (ii) for each particle location the vector P_normal_ was calculated, which represents the normal of S_estimate_ at that position (Figure 3 D); (iii) a vector P_denat_ was computed that describes the relative orientation of the denatured side as the vector pointing from the center of R_intact_ to the center of mass of R_missing_; (iv) the P_denat_ vector was calculated for every particle detected in all the tomographic volumes; (v) finally, the displacement angle *δ* between the vectors P_denat_ and P_normal_ was calculated, and the distribution of *δ* as a bild file (Figure 3 E) was plotted.

## Acknowledgements

We thank Deryck J. Mills, Simone Prinz and Mark Linder for EM support. We are grateful to Martin Centola, Niklas Klusch and Dr. David Wöhlert for discussions. We thank Dr. Janet Vonck and Dr. Roberto Covino for critically reading the manuscript. This project was funded by the Max Planck Society a Lichtenberg grant of the Volkswagen Foundation to M.G. (grant number 85701). This project was further supported by the LOEWE program (Landes-Offensive zur Entwicklung wissenschaftlichökonomischer Exzellenz) of the state of Hesse and was conducted within the framework of the MegaSyn Research Cluster.

## Author contributions statement

E.D.I. conceived and supervised the project. E.D.I. conducted all negative stain and single particle cryo-EM experiments.D.F. conducted all the cryo-ET and STA experiments. R.S. contributed to the orientational analysis of the particles in the tomographic volumes. M. J. performed the expression, purification, TSA and activity test assay of FAS complex. E.D.I. and D.F. analyzed the results. M.G and W.K. provided funds. All of the authors have taken part in the preparation of this manuscript, have reviewed the results, and have approved the final version of this manuscript.

## Additional information

The EM maps have been deposited in the EMDB with accession codes EMD-0178 (single particle cryo-EM FAS map on graphene support) and EMD-0179 (subtomogram averaging FAS map on graphene support).

**Figure 1 Supplement 1.**
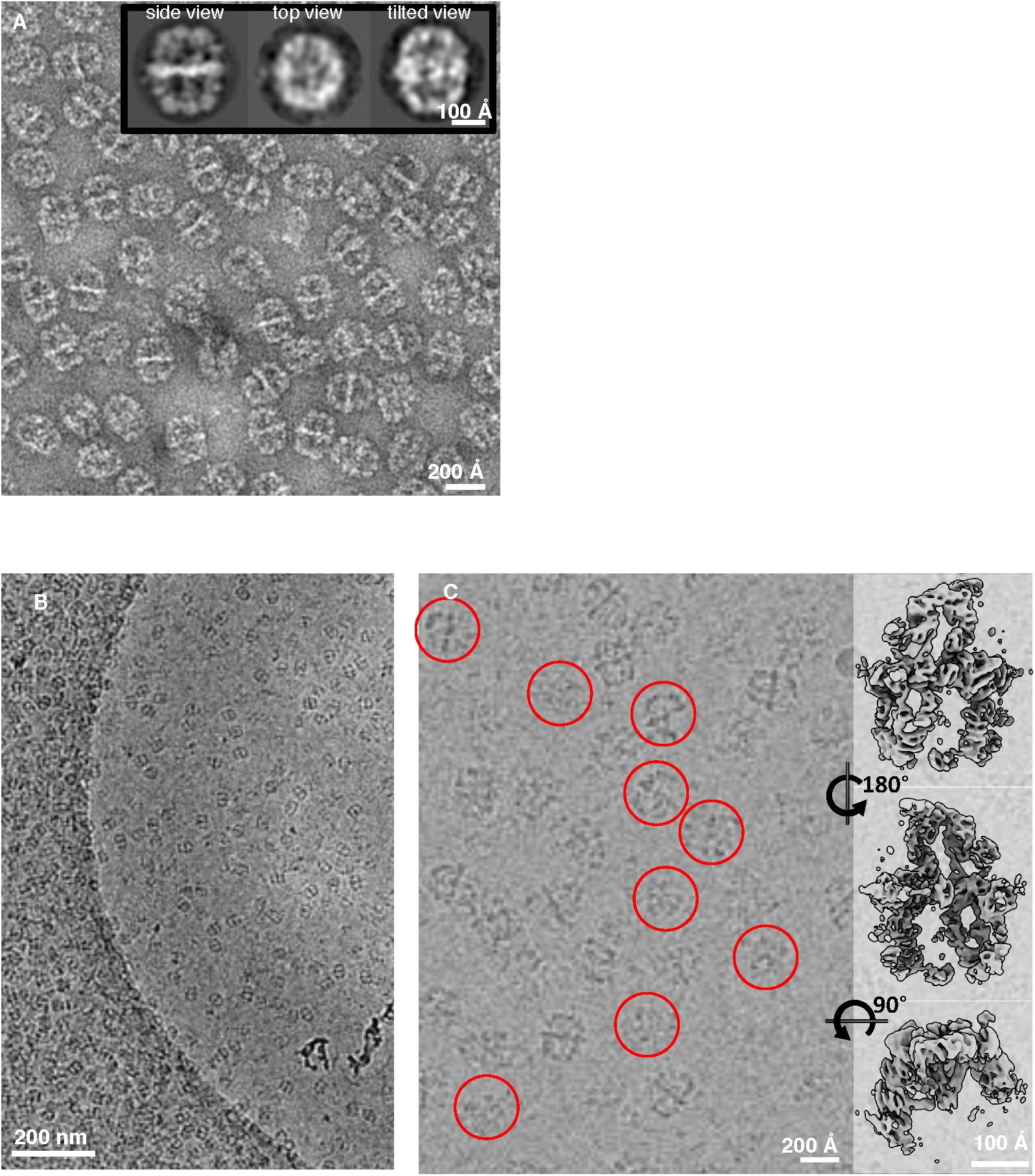
Negative stain versus standard cryo-EM specimens. (A) FAS (0.05 mg/ml) negatively stained with sodium silicotungstate. The inset shows three representative 2D class averages calculated with a reference-free classification. (B) Low magnification (8,700 x) overview of a Quantifoil R2/2 cryo grid hole. (C) Higher magnification (135,000x) image of an adjacent hole reveals some of the broken FAS particles (red circles), with three surface views of the 3D reconstruction from class 3 (Figure 1 B) on the right.

**Figure 1 Supplement 2.**
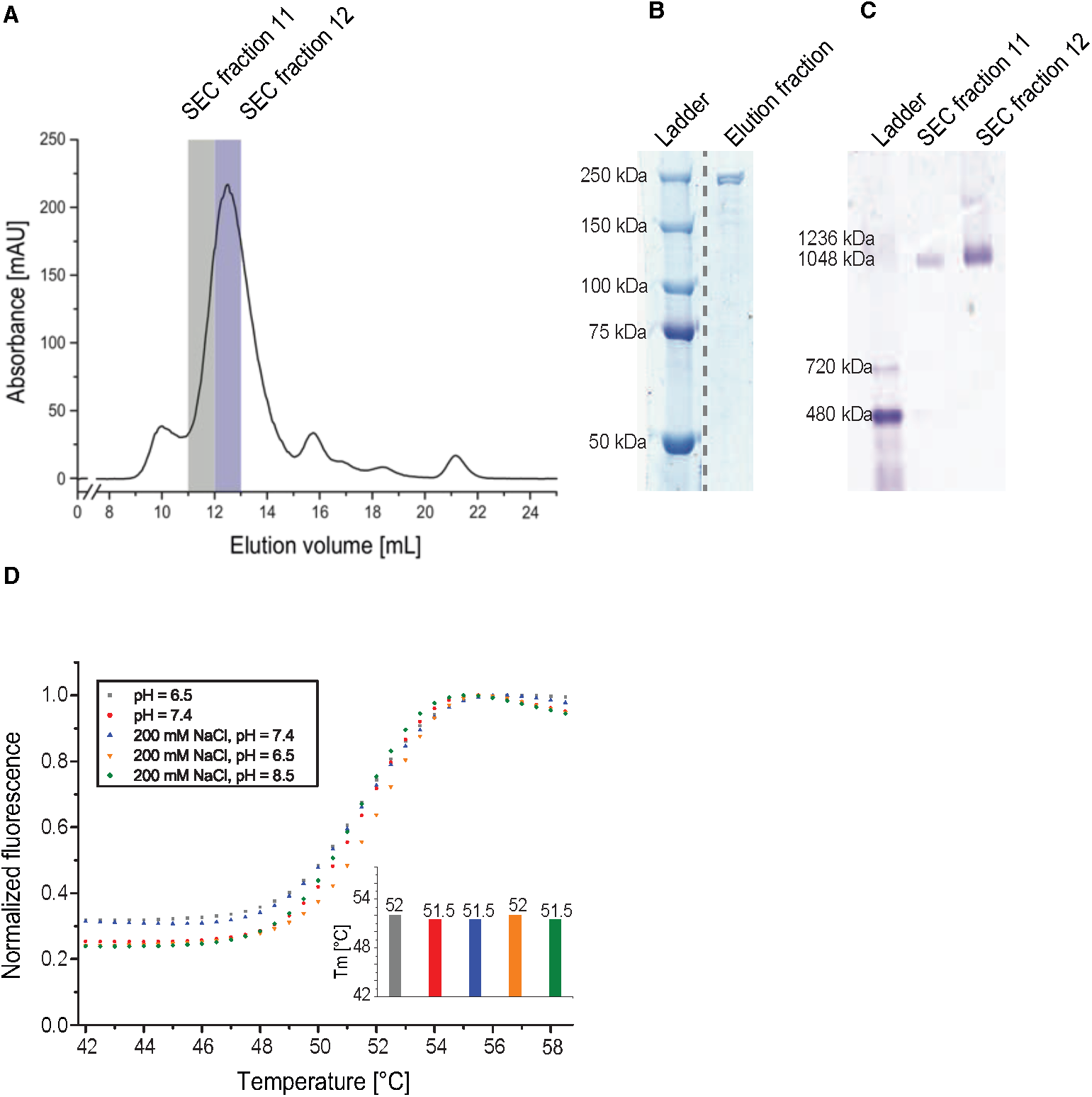
Purification and quality control of yeast FAS preparation. (A) Size exclusion chromatography (SEC) profile (Superose 6 Increase 10/300 GL, buffer: 100 mM sodium phosphate pH 6.5, fraction size: 1 mL. UV absorbance at 280 nm). (B) SDS PAGE of FAS affinity chromatography elution fraction (Ladder: Precision Plus Protein^TM^ Dual Color Standards, Bio rad). (C) Blue-native PAGE of SEC fractions (Ladder: NativeMark^TM^ Unstained Protein Ladder, Invitrogen^TM^). (D) Normalized thermal shift assays of yeast FAS in different buffers as a function of temperature. Columns show the average melting temperature of three replicates. The error of replicates was smaller than 0.5°C for each buffer, which is the limit of accuracy.

**Figure 2 Supplement 1.**
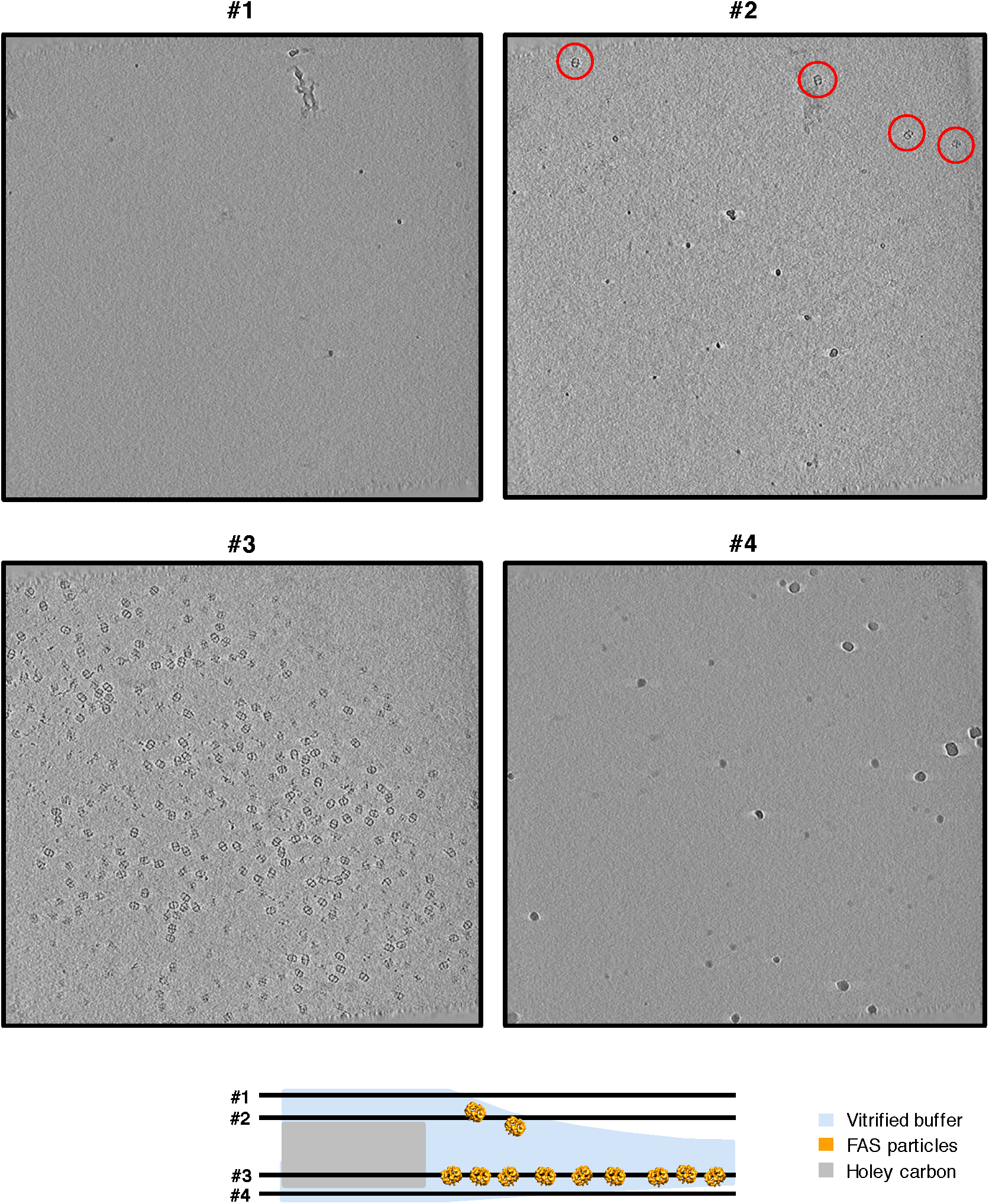
Slices through tomograms of unsupported vitrified solution. Four-pixel slices through NAD-filtered tomograms at various heights highlight atmospheric ice contamination (slices #1 and #4) used to identify the two air-water interfaces. The upper (FAS particles circled in red, slice #2) and lower (slice #3) meniscus of the vitrified buffer differ in the number of FAS particles, with the former being less populated.

**Figure 3 Supplement 1.**
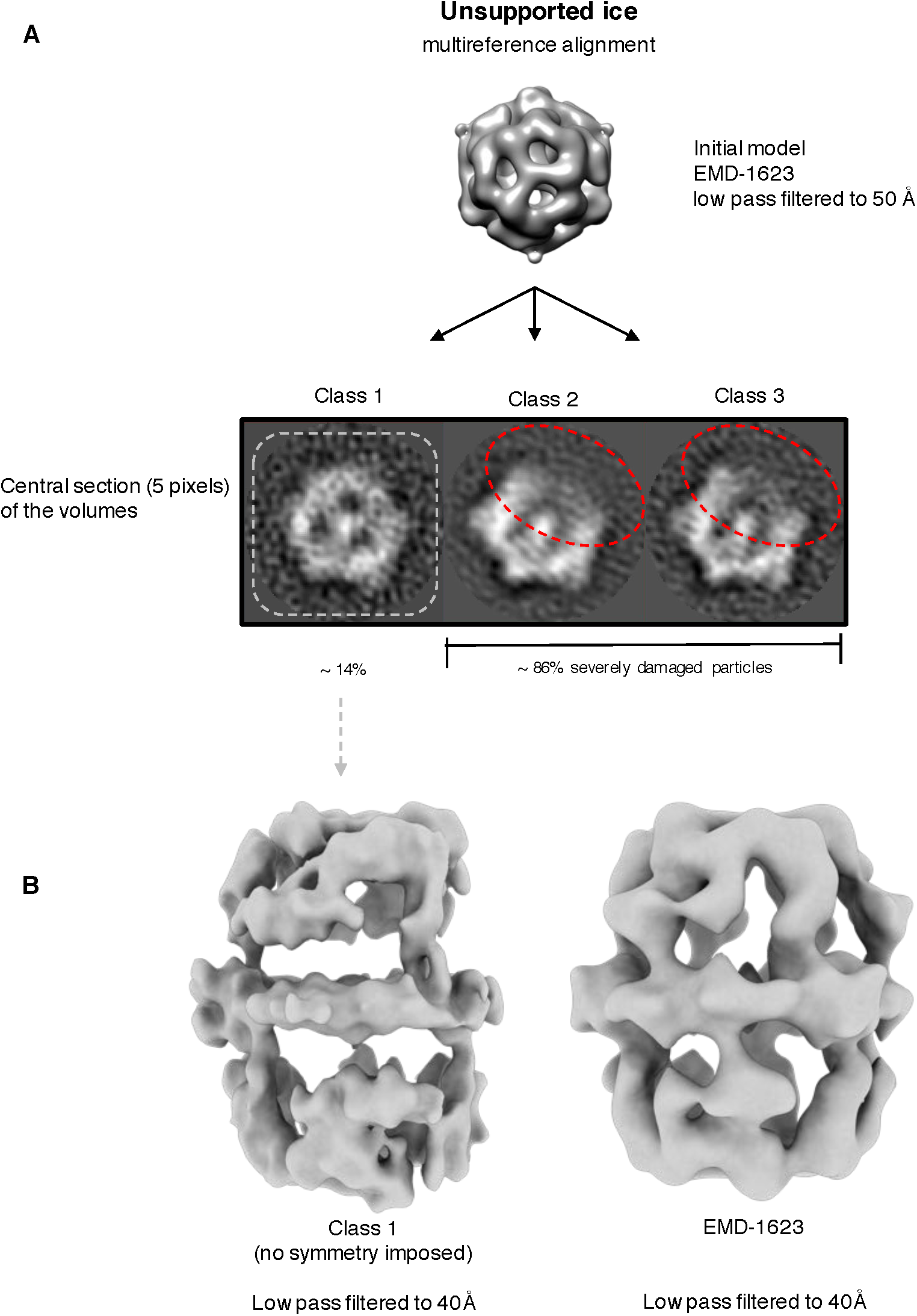
Multireference alignment of FAS in unsupported vitrified solution. (A) Classification by MRA indicates FAS denaturation comparable to the single-particle results (Figure 1 B). The particles reconstructed from the best class (class 1) led to a map with signs of denaturation (B). A map of intact FAS (EMD-1623) is shown for comparison.

**Figure 6 Supplement 1.**
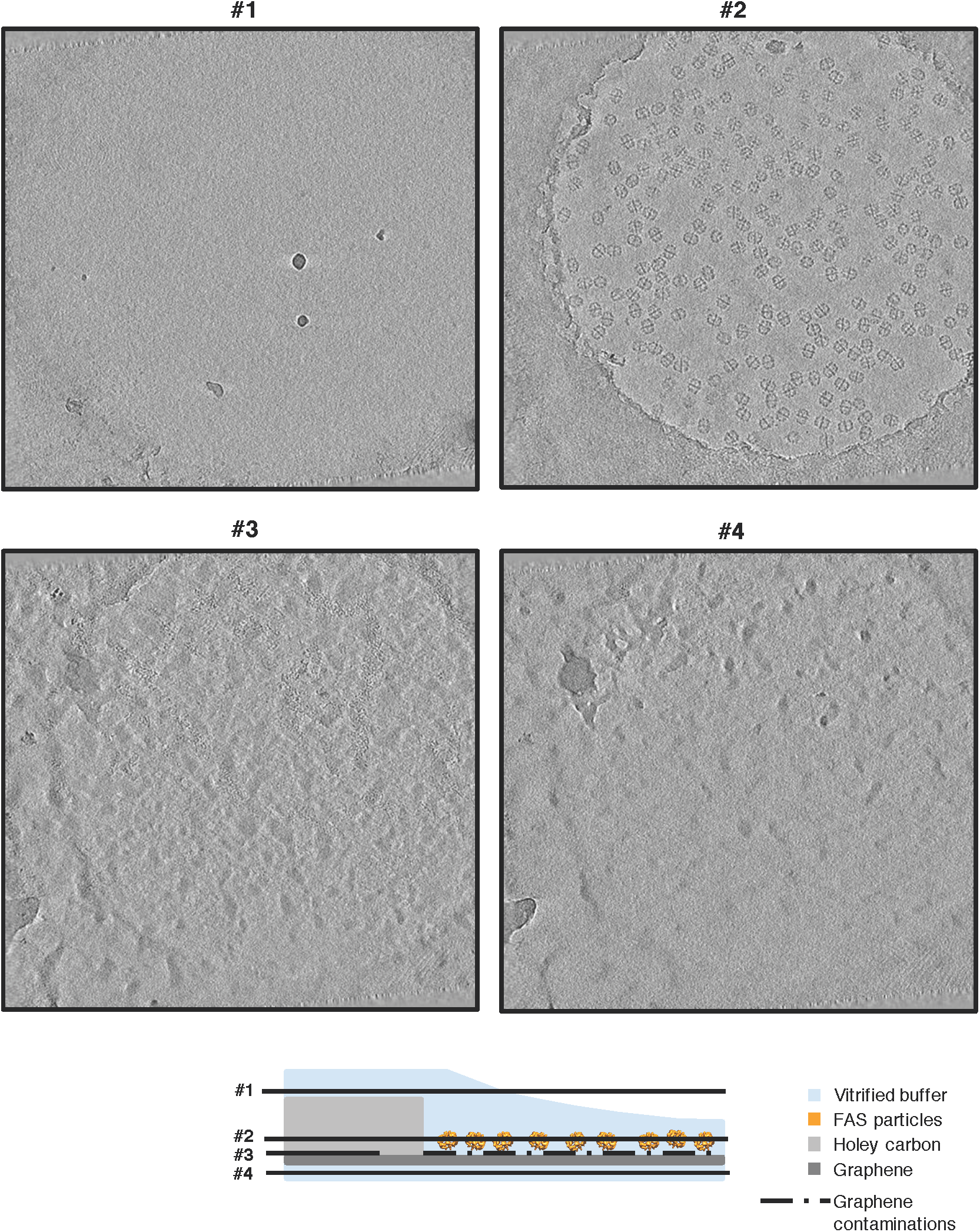
Slices through the tomograms from specimen with hydrophilized graphene. Four-pixel slices through NAD-filtered tomograms at various heights show contaminating ice crystals (slices #1 and #4) used to identify the two air-water interfaces. FAS particles are evenly spread in a single layer (slice #2) on the graphene film. A layer of contaminants on the lower surface identifies the position of the electron transparent graphene sheet (slice #3).

**Figure 6 Supplement 2.**
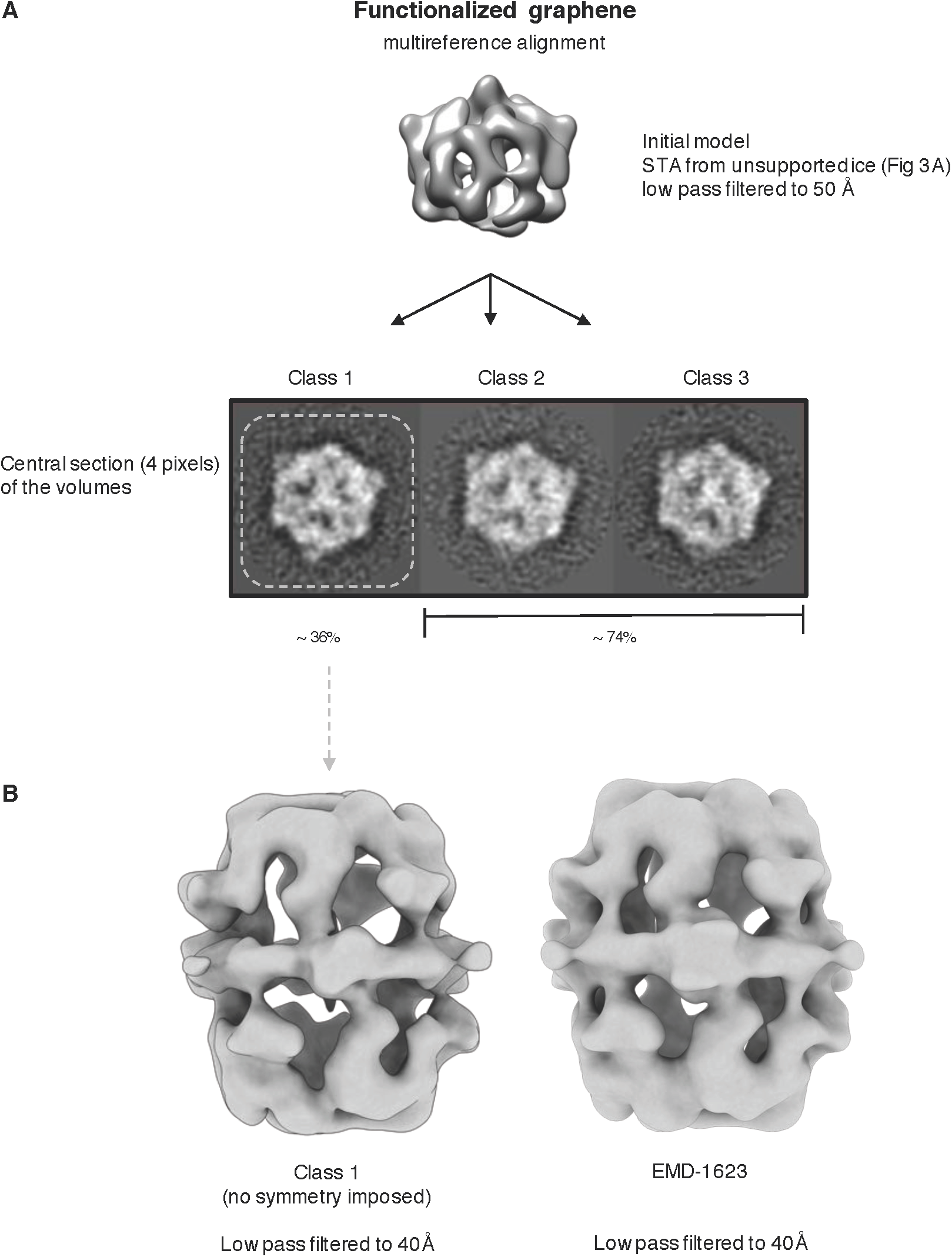
FAS particles in vitrified buffer on hydrophilized graphene. Classification by MRA (A) indicates that all particles on hydrophilized graphene support are intact, even though the average volume of partly denatured FAS (Figure 3 A) was used as an initial reference to avoid bias. The map resulting from the best class (1) does not show any sign of denaturation (B). A map of intact FAS (EMD-1623) is included for comparison.

**Figure 6 Supplement 3.**
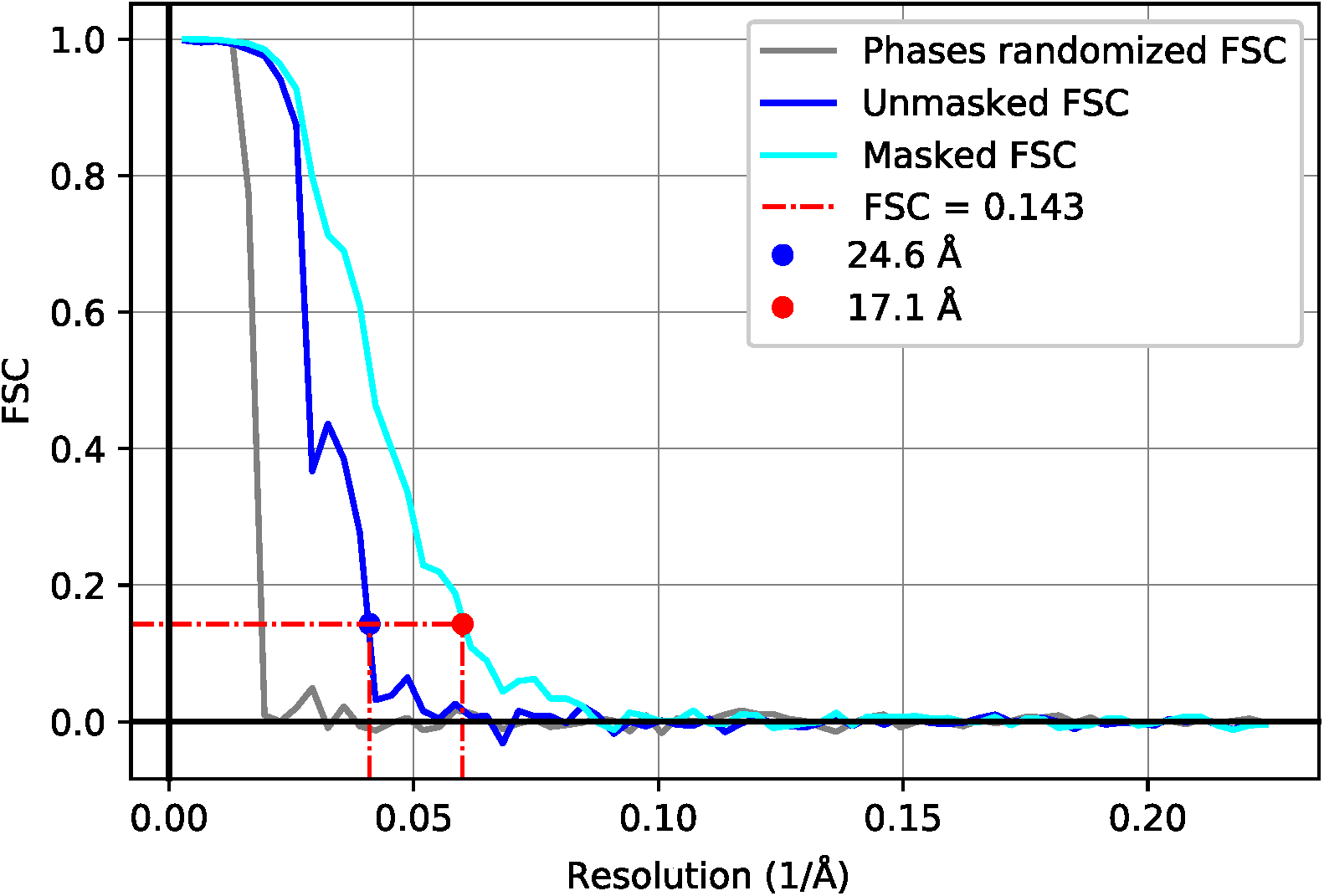
Resolution estimation of subtomogram averages. FSC of the two unfiltered half-maps of FAS complex before (blue) and after (cyan) masking. FSC performed on the half-maps with phases randomized beyond 60 Å (grey). FSC curves before and after masking indicate resolutions of 24.6 Å and 17.1 Å respectively.

**Figure 8 Supplement 1.**
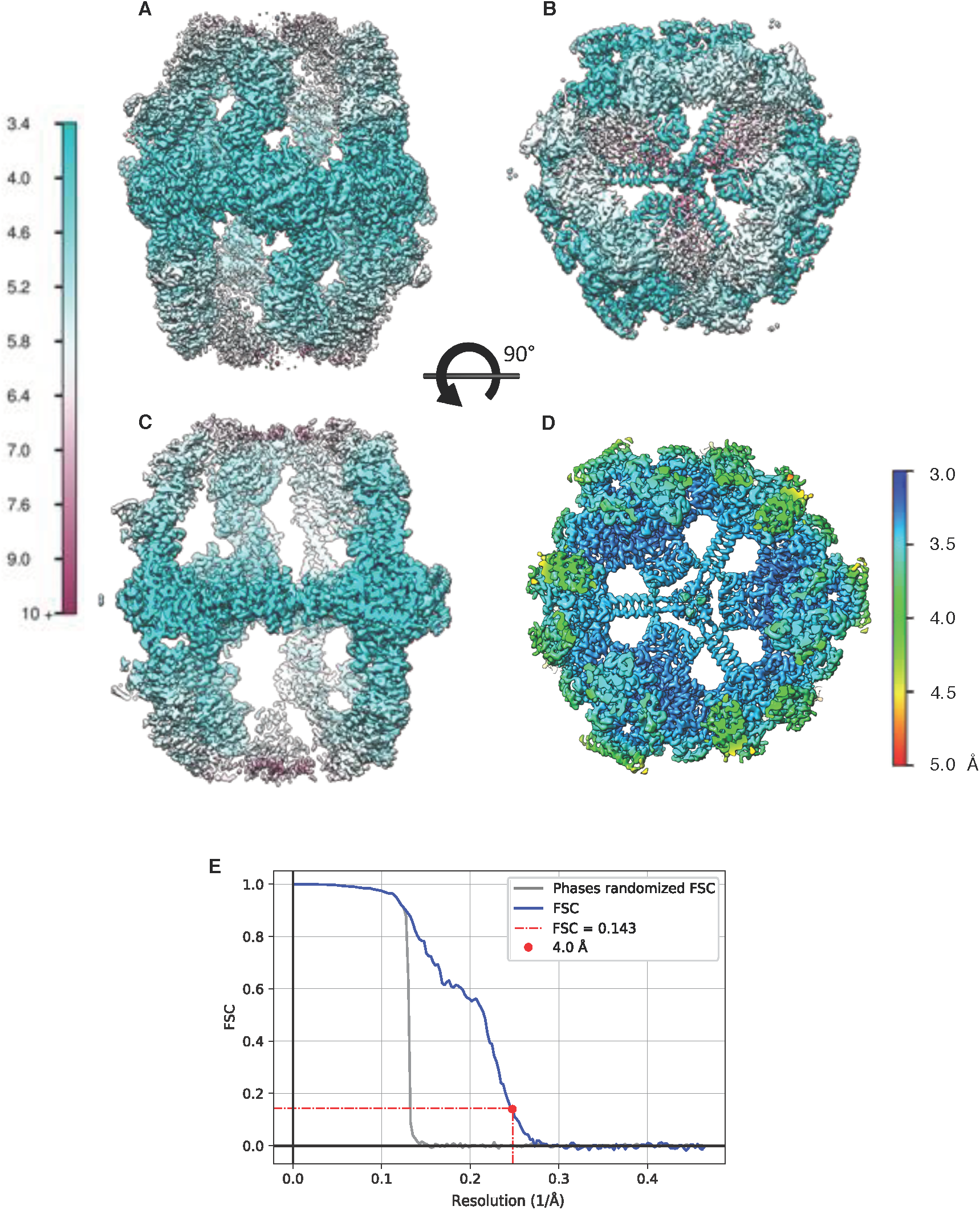
Local resolution estimates. Local resolution estimation of FAS complex refined to a global resolution of 4.0 Å (Figure 8 D) as seen in side view (A) and top view (B). (C) Longitudinal section of (A). (D) Transversal section of (B) with a local map resolution beyond 3.5 Å. (E) Fourier shell correlation of the two half-maps, unfiltered (blue) and phase-randomized (grey).

**Figure 8 Supplement 2.**
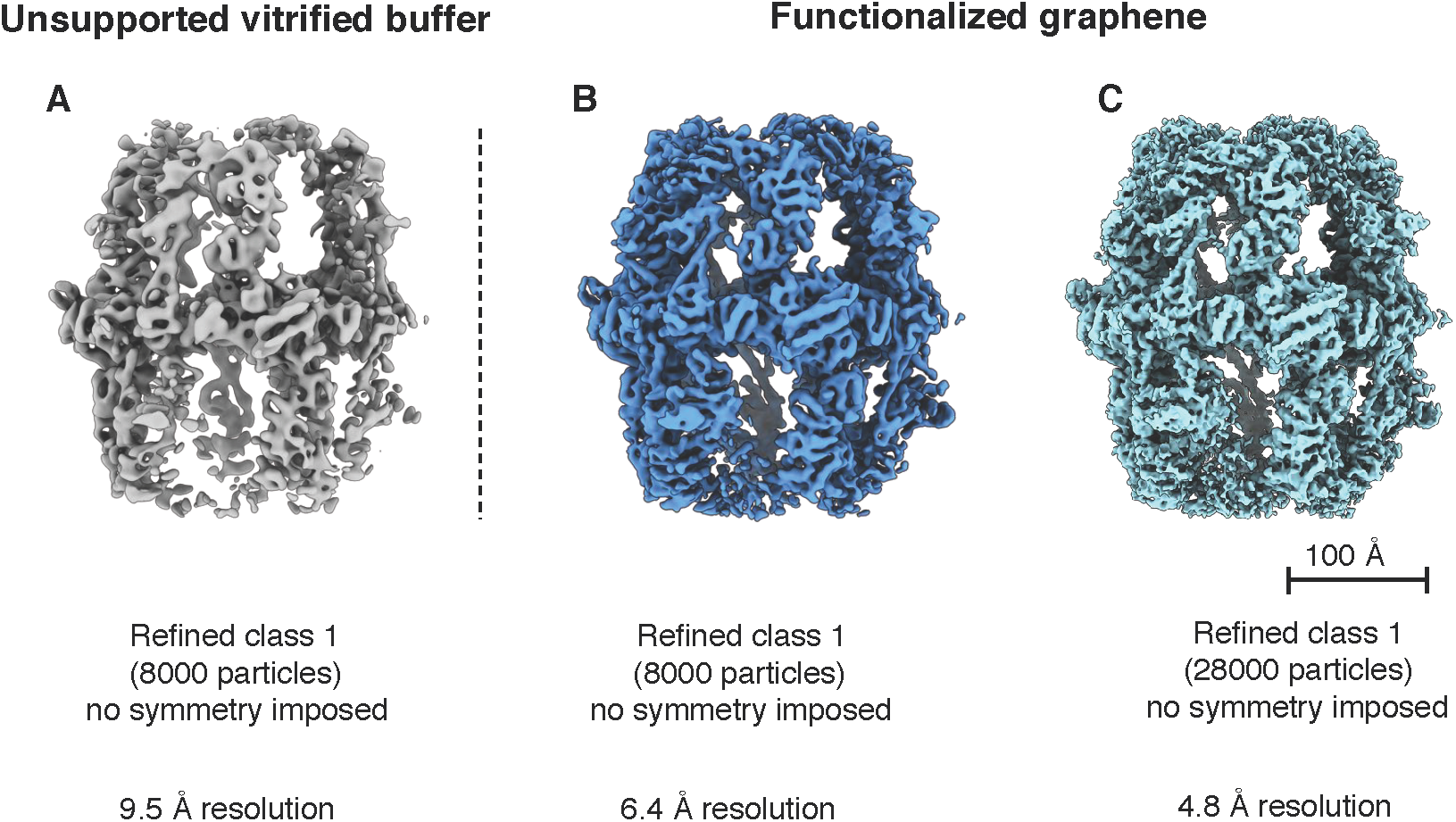
Reconstructions from different particle numbers. (A) Unsymmetrized reconstruction of FAS complex from 8,000 particle images in unsupported vitrified buffer (Class 1 from Figure 1 B) compared with the map of FAS on hydrophilized graphene (Class 1 from Figure 8 B) from the same number of randomly selected particles (B). (C) The best asymmetric reconstruction of FAS complex from 28,000 particles (Class 1 from Figure 8 B).

**Figure 8 Supplement 3.**
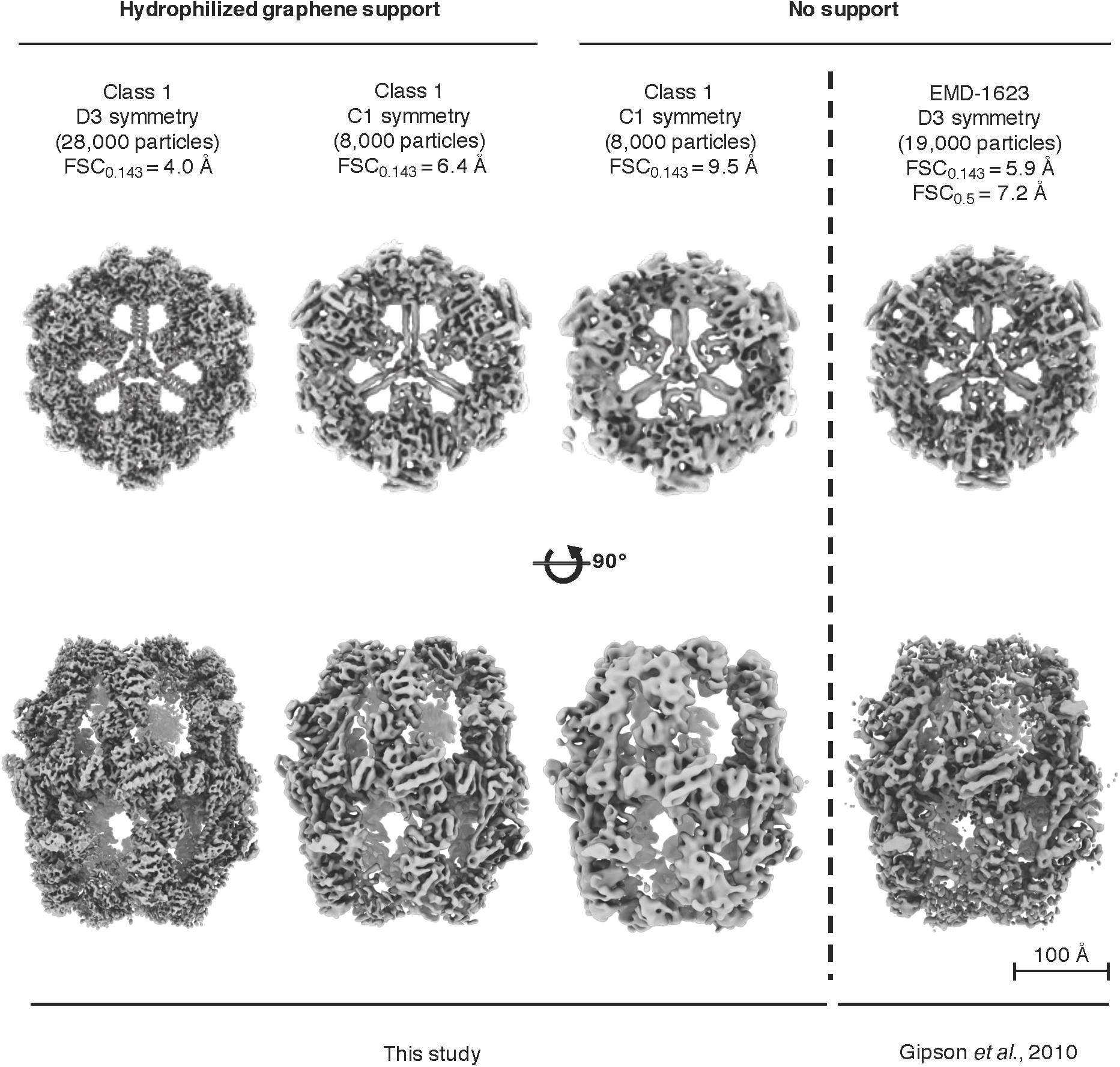
Differences in the quality of map reconstruction. Comparison between Cryo-EM maps reconstructed in this study and the map reported in Gipson et al., 2010^39^. The quality of the EMD-1623 map is intermediate between our FAS reconstruction in unsupported vitrified buffer, and the one obtained in the same conditions with hydrophilized graphene support.

## References

1. Kühlbrandt, W. The resolution revolution. Sci. 343, 1443–1444 (2014). DOI 10.1126/science.1251652.

2. Bai, X.-c., McMullan, G. & Scheres, S. H. W. How cryo-em is revolutionizing structural biology. Trends Biochem. Sci. 40, 49–57 (2015). URL. DOI http://dx.doi.org/10.1016/j.tibs.2014.10.005.

3. Cheng, Y., Grigorieff, N., Penczek, P. A. & Walz, T. A primer to single-particle cryo-electron microscopy. Cell 161, 438–49 (2015). URL. DOI 10.1016/j.cell.2015.03.050.

4. Passmore, L. A. & Russo, C. J. Specimen preparation for high-resolution cryo-em. Methods Enzym. 579, 51–86 (2016). URL. DOI 10.1016/bs.mie.2016.04.011.

5. McDowall, A. et al. Electron microscopy of frozen hydrated sections of vitreous ice and vitrified biological samples. J. Microsc. 131, 1–9 (1983). URL. DOI 10.1111/j.1365-2818.1983.tb04225.x.

6. Dubochet, J. et al. Cryo-electron microscopy of vitrified specimens. Q. Rev. Biophys. 21, 129–228 (1988). URL. DOI 10.1017/S0033583500004297.

7. Ramsden, J. J. Experimental methods for investigating protein adsorption kinetics at surfaces. Q Rev Biophys 27, 41–105 (1994). URL. DOI 10.1017/S0033583500002900.

8. Trurnit, H. A theory and method for the spreading of protein monolayers. J. Colloid Sci. 15, 1–13 (1960). URL. DOI 10.1016/0095-8522(60)90002-7.

9. Neurath, H. & Bull, H. B. The surface activity of proteins. Chem. Rev. 23, 391–435 (1938).URL. DOI 10.1021/cr60076a001.

10. Noble, A. J. et al. Routine single particle cryoem sample and grid characterization by tomography. Elife 7 (2018). URL. DOI 10.7554/eLife.34257.

11. Israelachvili, J. N. Intermolecular and surface forces (Academic press, 2011). URL.

12. Naydenova, K. & Russo, C. J. Measuring the effects of particle orientation to improve the efficiency of electron cryomicroscopy. Nat Commun 8, 629 (2017). URL. DOI 10.1038/s41467-017-00782-3.

13. Jain, T., Sheehan, P., Crum, J., Carragher, B. & Potter, C. S. Spotiton: a prototype for an integrated inkjet dispense and vitrification system for cryo-tem. J Struct Biol 179, 68–75 (2012). URL. DOI 10.1016/j.jsb.2012.04.020.

14. Wei, H. et al. Optimizing “self-wicking” nanowire grids. J Struct Biol 202, 170–174 (2018).URL. DOI 10.1016/j.jsb.2018.01.001.

15. Dandey, V. P. et al. Spotiton: New features and applications. J Struct Biol 202, 161–169 (2018). URL. DOI 10.1016/j.jsb.2018.01.002.

16. Scapin, G. et al. Structure of the insulin receptor-insulin complex by single-particle cryo-em analysis. Nat. 556, 122–125 (2018). URL. DOI 10.1038/nature26153.

17. Xu, K. et al. Epitope-based vaccine design yields fusion peptide-directed antibodies that neutralize diverse strains of hiv-1. Nat Med 24, 857–867 (2018). URL. DOI 10.1038/s41591-018-0042-6.

18. Popot, J. L. Amphipols, nanodiscs, and fluorinated surfactants: three nonconventional approaches to studying membrane proteins in aqueous solutions. Annu. Rev Biochem. 79, 737–75 (2010). URL. DOI 10.1146/annurev.biochem.052208.114057.

19. Blees, A. et al. Structure of the human mhci peptide-loading complex. Nat. 551, 525–528 (2017). URL. DOI 10.1038/nature24627.

20. Efremov, R. G., Leitner, A., Aebersold, R. & Raunser, S. Architecture and conformational switch mechanism of the ryanodine receptor. Nat. 517, 39–43 (2015). URL. DOI 10.1038/nature13916.

21. Bai, X.-c., Fernandez, I. S., McMullan, G. & Scheres, S. H. Ribosome structures to near-atomic resolution from thirty thousand cryo-em particles. eLife 2, e00461 (2013). URL. DOI 10.7554/eLife.00461.

22. D’Imprima, E. et al. Cryoem structure of the bifunctional secretin complex of thermus thermophilus. Elife 6 (2017). URL. DOI 10.7554/eLife.30483.

23. Nguyen, T. H. et al. The architecture of the spliceosomal u4/u6.u5 tri-snrnp. Nat. 523, 47–52 (2015). URL. DOI 10.1038/nature14548.

24. Schraidt, O. & Marlovits, T. C. Three-dimensional model of salmonella’s needle complex at subnanometer resolution. Sci. 331, 1192–5 (2011). URL. DOI 10.1126/science.1199358.

25. Brink, J., Sherman, M., Berriman, J. & Chiu, W. Evaluation of charging on macromolecules in electron cryomicroscopy. Ultramicroscopy 72, 41–52 (1998). URL. DOI 10.1016/S0304-3991(97)00126-5.

26. Larson, D. M., Downing, K. H. & Glaeser, R. M. The surface of evaporated carbon films is an insulating, high-bandgap material. J Struct Biol 174, 420–3 (2011). URL. DOI 10.1016/j.jsb.2011.02.005.

27. Russo, C. J. & Henderson, R. Charge accumulation in electron cryomicroscopy. Ultramicroscopy 187, 43–49 (2018). URL. DOI 10.1016/j.ultramic.2018.01.009.

28. Russo, C. J. & Henderson, R. Microscopic charge fluctuations cause minimal contrast loss in cryoem. Ultramicroscopy 187, 56–63 (2018). URL. DOI 10.1016/j.ultramic.2018.01.011.

29. Russo, C. J. & Passmore, L. A. Controlling protein adsorption on graphene for cryo-em using low-energy hydrogen plasmas. Nat Methods 11, 649–52 (2014).URL. DOI 10.1038/nmeth.2931.

30. Geim, A. K. & Novoselov, K. S. The rise of graphene. Nat Mater 6, 183–91 (2007). URL. DOI 10.1038/nmat1849.

31. Sader, K., Stopps, M., Calder, L. J. & Rosenthal, P. B. Cryomicroscopy of radiation sensitive specimens on unmodified graphene sheets: reduction of electron-optical effects of charging. J Struct Biol 183, 531–536 (2013). URL. DOI 10.1016/j.jsb.2013.04.014.

32. Pantelic, R. S., Meyer, J. C., Kaiser, U. & Stahlberg, H. The application of graphene as a sample support in transmission electron microscopy. Solid State Commun. 152, 1375–1382 (2012). URL. DOI http://dx.doi.org/10.1016/j.ssc.2012.04.038.

33. Pantelic, R. S., Meyer, J. C., Kaiser, U., Baumeister, W. & Plitzko, J. M. Graphene oxide: a substrate for optimizing preparations of frozen-hydrated samples. J Struct Biol 170, 152–6 (2010). URL. DOI 10.1016/j.jsb.2009.12.020.

34. Boland, A. et al. Cryo-em structure of a metazoan separase-securin complex at near-atomic resolution. Nat Struct Mol Biol 24, 414–418 (2017). URL. DOI 10.1038/nsmb.3386.

35. Pantelic, R. S., Fu, W., Schoenenberger, C. & Stahlberg, H. Rendering graphene supports hydrophilic with non-covalent aromatic functionalization for transmission electron microscopy. Appl. Phys. Lett. 104, 134103 (2014). URL. DOI 10.1063/1.4870531.

36. Jenni, S. et al. structure of fungal fatty acid synthase and implications for iterative substrate shuttling. Sci. 316, 254–261 (2007). URL. DOI 10.1126/science.1138248.

37. Johansson, P. et al. Inhibition of the fungal fatty acid synthase type i multienzyme complex. Proc. Natl. Acad. Sci. USA 105, 12803–12808 (2008). URL. DOI 10.1073/pnas.0805827105.

38. Lomakin, I., Xiong, Y. & Steitz, T. The crystal structure of yeast fatty acid synthase, a cellular machine with eight active sites working together. Cell 129, 319–332 (2007). URL. DOI 10.1016/j.cell.2007.03.013.

39. Gipson, P. et al. Direct structural insight into the substrate shuttling mechanism of yeast fatty acid synthase by electron cryo-microscopy. Proc. Natl. Acad. Sci. USA 107, 9164–9169 (2010). DOI 10.1073/pnas.0913547107.

40. Fichtlscherer, F., Wellein, C., Mittag, M. & Schweizer, E. A novel function of yeast fatty acid synthase. subunit alpha is capable of self-pantetheinylation. Eur J Biochem. 267, 2666–71 (2000). URL.

41. Oesterhelt, D., Bauer, H. & Lynen, F. Crystallization of a multienzyme complex: fatty acid synthetase from yeast. Proc Natl Acad Sci U S A 63, 1377–82 (1969). URL. DOI 10.1073/pnas.63.4.1377.

42. Wieland, F., Renner, L., Verfurth, C. & Lynen, F. Studies on the multi-enzyme complex of yeast fatty-acid synthetase. reversible dissociation and isolation of two polypeptide chains. Eur J Biochem. 94, 189–97 (1979). URL. DOI 10.1111/j.1432-1033.1979.tb12885.x.

43. Scheres, S. H. & Chen, S. Prevention of overfitting in cryo-em structure determination. Nat Methods 9, 853–4 (2012). URL. DOI 10.1038/nmeth.2115.

44. Chen, S. et al. High-resolution noise substitution to measure overfitting and validate resolution in 3d structure determination by single particle electron cryomicroscopy. Ultramicroscopy 135, 24–35 (2013). URL. DOI 10.1016/j.ultramic.2013.06.004.

45. Kastritis, P. L. et al. Capturing protein communities by structural proteomics in a thermophilic eukaryote. Mol Syst Biol 13, 936 (2017). URL. DOI 10.15252/msb.20167412.

46. Boehringer, D., Ban, N. & Leibundgut, M. 7.5-Å cryo-em structure of the mycobacterial fatty acid synthase. J. Mol. Biol. 425, 841–849 (2013). DOI 10.1016/j.jmb.2012.12.021.

47. Zhu, S. et al. Structure of a human synaptic gabaa receptor. Nat. 559, 67–72 (2018). URL. DOI 10.1038/s41586-018-0255-3.

48. Kim, Y. & Chen, J. Molecular structure of human p-glycoprotein in the atp-bound, outward-facing conformation. Sci. 359, 915–919 (2018). URL. DOI 10.1126/science.aar7389.

49. Takizawa, Y. et al. Cryoem structure of the nucleosome containing the alb1 enhancer dna sequence. Open Biol 8 (2018). URL. DOI 10.1098/rsob.170255.

50. Bai, X. C. et al. An atomic structure of human gamma-secretase. Nat. 525, 212–7 (2015). URL. DOI 10.1038/nature14892.

51. Yan, Z. et al. Structure of the nav1.4-beta1 complex from electric eel. Cell 170, 470–482 e11 (2017). URL. DOI 10.1016/j.cell.2017.06.039.

52. Chakravarty, B., Gu, Z., Chirala, S., Wakil, S. & Quiocho, F. Human fatty acid synthase: structure and substrate selectivity of the thioesterase domain. Proc. Natl. Acad. Sci. USA 101, 15567–15572 (2004).

53. Gajewski, J., Pavlovic, R., Fischer, M., Boles, E. & Grininger, M. Engineering fungal de novo fatty acid synthesis for short chain fatty acid production. Nat Commun 8, 14650 (2017). URL. DOI 10.1038/ncomms14650.

54. Ericsson, U. B., Hallberg, B. M., Detitta, G. T., Dekker, N. & Nordlund, P. Thermofluor-based high-throughput stability optimization of proteins for structural studies. Anal Biochem. 357, 289–98 (2006). URL. DOI 10.1016/j.ab.2006.07.027.

55. Gajewski, J. et al. Engineering fatty acid synthases for directed polyketide production. Nat Chem Biol 13, 363–365 (2017). URL. DOI 10.1038/nchembio.2314.

56. Grant, T. & Grigorieff, N. Measuring the optimal exposure for single particle cryo-em using a 2.6 Å reconstruction of rotavirus vp6. eLife 4, e06980 (2015). URL. DOI 10.7554/eLife.06980.

57. Zheng, S. Q. et al. Motioncor2: anisotropic correction of beam-induced motion for improved cryo-electron microscopy. Nat Methods 14, 331–332 (2017). URL. DOI 10.1038/nmeth.4193.

58. Rohou, A. & Grigorieff, N. Ctffind4: Fast and accurate defocus estimation from electron micrographs. J Struct Biol 192, 216–21 (2015). URL. DOI 10.1016/j.jsb.2015.08.008.

59. McMullan, G., Vinothkumar, K. R. & Henderson, R. Thon rings from amorphous ice and implications of beam-induced brownian motion in single particle electron cryo-microscopy. Ultramicroscopy 158, 26–32 (2015). URL. DOI 10.1016/j.ultramic.2015.05.017.

60. Kimanius, D., Forsberg, B. O., Scheres, S. H. & Lindahl, E. Accelerated cryo-em structure determination with parallelisation using gpus in relion-2. Elife 5 (2016). URL. DOI 10.7554/eLife.18722.

61. Goddard, T. D. et al. Ucsf chimerax: Meeting modern challenges in visualization and analysis. Protein Sci 27, 14–25 (2018). URL. DOI 10.1002/pro.3235.

62. Hagen, W. J. H., Wan, W. & Briggs, J. A. G. Implementation of a cryo-electron tomography tilt-scheme optimized for high resolution subtomogram averaging. J Struct Biol 197, 191–198 (2017). URL. DOI 10.1016/j.jsb.2016.06.007.

63. Mastronarde, D. N. Automated electron microscope tomography using robust prediction of specimen movements. J Struct Biol 152, 36–51 (2005). URL. DOI 10.1016/j.jsb.2005.07.007.

64. Zhang, K. Gctf: realtime ctf determination and correction. J. Struct. Biol. 193, 1–12 (2016).URL. DOI 10.1016/j.jsb.2015.11.003.

65. Kremer, J. R., Mastronarde, D. N. & McIntosh, J. R. Computer visualization of three-dimensional image data using imod. J Struct Biol 116, 71–6 (1996). URL. DOI 10.1006/jsbi.1996.0013.

66. Frangakis, A. S. & Hegerl, R. Noise reduction in electron tomographic reconstructions using nonlinear anisotropic diffusion. J Struct Biol 135, 239–50 (2001). URL. DOI 10.1006/jsbi.2001.4406.

67. Castano-Diez, D. The dynamo package for tomography and subtomogram averaging: components for matlab, gpu computing and ec2 amazon web services. Acta Crystallogr D Struct Biol 73, 478–487 (2017). URL. DOI 10.1107/S2059798317003369.

68. Pettersen, E. F. et al. Ucsf chimera–a visualization system for exploratory research and analysis. J Comput. Chem 25, 1605–12 (2004). URL. DOI 10.1002/jcc.20084.

69. Chen, M. et al. Convolutional neural networks for automated annotation of cellular cryo-electron tomograms. Nat Methods 14, 983–985 (2017). URL. DOI 10.1038/nmeth.4405.

70. Tang, G. et al. Eman2: An extensible image processing suite for electron microscopy. J. Struct. Biol. 157, 38–46 (2007). DOI 10.1016/j.jsb.2006.05.009.

